# Armadillo Repeat Only Proteins Are Essential Activators of Plant CNGC Channels

**DOI:** 10.1101/2025.01.06.631460

**Authors:** Ivan Kulich, Denisa Oulehlová, Dmitrii Vladimirtsev, Samuel Boscq, Lukáš Martínek, Minxia Zou, Alexey Bondar, Katarína Kulichová, Edita Lileikyte, Serban Pop, Martin Janda, Oksana Iakovenko, Michaela Neubergerová, Armin Uebel, Florian Rauch, Tanja Studtrucker, Roman Pleskot, Petra Dietrich, Matyáš Fendrych, Jiří Friml

## Abstract

The versatile Ca^2+^ signaling system governs plant responses to a wide array of environmental and developmental cues. CYCLIC NUCLEOTIDE-GATED CHANNELS (CNGCs) trigger cellular responses to diverse signals, including phytohormones, biotic and abiotic stresses; and as such their activity is tightly controlled. Unlike their animal paralogs, plant CNGCs does not seem to be gated by cyclic nucleotides, and the mechanism of their activation remains unresolved. Here we report ARMADILLO REPEAT ONLY (ARO) proteins as novel, plant-specific, and essential activators of plant CNGCs. Reciprocal proximity labeling revealed interactions between all sporophytic CNGCs and AROs. Loss-of-function *aro* mutants fail to induce Ca^2+^ transients in response to known CNGC-triggering stimuli. Structural modeling, mutational analysis, and electrophysiological data show that AROs assemble into a complex with CNGC tetramers by interaction with the conserved Calmodulin-binding IQ domain. AROs represent CNGC activators, competing with Calmodulins, showcasing an evolutionarily unique solution to regulation of calcium signaling in plants.

## Introduction

Signaling mediated by cytoplasmic Ca^2+^ is an indispensable mechanism conserved across eukaryotes. Plants use Ca^2+^ signaling to respond to external and internal stimuli. CYCLIC NUCLEOTIDE GATED CHANNELS (CNGCs) are a class of tetrameric ion channels, which are essential for a broad spectrum of processes including hormonal responses^1–3^, stomatal movements^4,5^, tip growth^6,7^, and responses to microbe and damage-associated molecular patterns (MAMPs, DAMPs, respectively^8–10^). Responses to specific stimuli usually require a specific channel, with CNGC2 being downstream of extracellular ATP (eATP^11–13^), brassinosteroids^1^, and plant elicitor peptides (PEP) response^14^; CNGC14 downstream of phytohormone auxin; CNGC12 downstream of flagellin peptide FLG22^15^ etc.

Although numerous signaling pathways and adaptation responses are known to involve CNGCs, their exact operational mechanisms remain unclear. In heterologous systems, CNGCs exhibit hyperpolarization-activated conductance, suggesting voltage gating. However, other studies indicate ligand gating, wherein activation by cAMP or cGMP may be voltage-dependent or independent^7,9,16–18^. Recent structural studies^19,20^ suggest that plant CNGCs are not gated by cyclic nucleotides, due to a mutation in the cyclic nucleotide binding domain. The mode of their activation in plants thus remains unclear. Calmodulin gating was also implied; calmodulins were shown as negative regulators of CNGC opening^21,22^. Multiple studies highlight phosphorylation as the mechanism of CNGC regulation *in planta*^5,15,23,24^, however, it is unclear whether and how this contributes to the gating mechanism. To summarize, there is a lack of clear consensus on the CNGC gating, which signifies a deeper lack of understanding of the CNGC biology.

In *Arabidopsis thaliana*, Armadillo Repeat Only (ARO) constitue a class of four proteins which contain two sets of ARM repeats separated by an unstructured linker region. Initially, ARO1 was described to be essential for pollen tube growth^16^. Sporophytic AROs (ARO2, 3 and 4) were shown to be essential for trichome and root hair growth in the Arabidopsis sporophyte^25^. As in the pollen tubes, *aro2/3/4* mutant root hairs either abort or burst soon after bulging. This has been linked to the inability to spatially restrict ROP GTPase activity and to the loss of polarity^25^. Importantly, all AROs are functionally redundant. ARO from *Marchantia polymorpha* complemented Arabidopsis *aro2/3/4* mutant phenotypes demonstrating a large degree of conservation of its function in land plants^25^.

Here we provide evidence that the major cause of the *aro2/3/4* mutant phenotypes is a lack of function of multiple CNGCs, and establish AROs as interactors and indispensable accessory proteins for CNGC function.

## Results

### AROs interact with CNGCs

As we showed previously, all four ARO isoforms from *Arabidopsis thaliana* and ARO from *Marchantia polymorpha* are functionally redundant^25^. We therefore focused on ARO2, which has the most general expression pattern^25^. To detect interactors of AROs, we performed co-immunoprecipitation and Mass Spectrometry with plants stably transformed with ARO2-HA-TurboID (tID). A polybasic stretch tagged with GFP (PBR-GFP-HA-tID) was used as a negative control to mimic AROs polybasic region^17^. We identified multiple CNGCs enriched significantly on ARO2 bait (Fig. S1). To increase sensitivity of the experiment, we then performed proximity labelling with ARO2-HA-tID. Of the 20 CNGCs present in Arabidopsis genome, 15 were identified in all replicates (two triplicates), while no signal was found in the negative control. Most (11) of the CNGCs were found in the top 30 most differentially biotinylated proteins in all replicates (Fig. 1A, B). Interestingly, the five remaining CNGCs absent in the ARO2 proxiome are not expressed in the seedling stage of Arabidopsis (Fig. S1B). Among enriched proteins, we also identified multiple IQ domain (IQD) proteins, which are a class of calmodulin binding microtubular proteins^26^.

**Figure 1.**
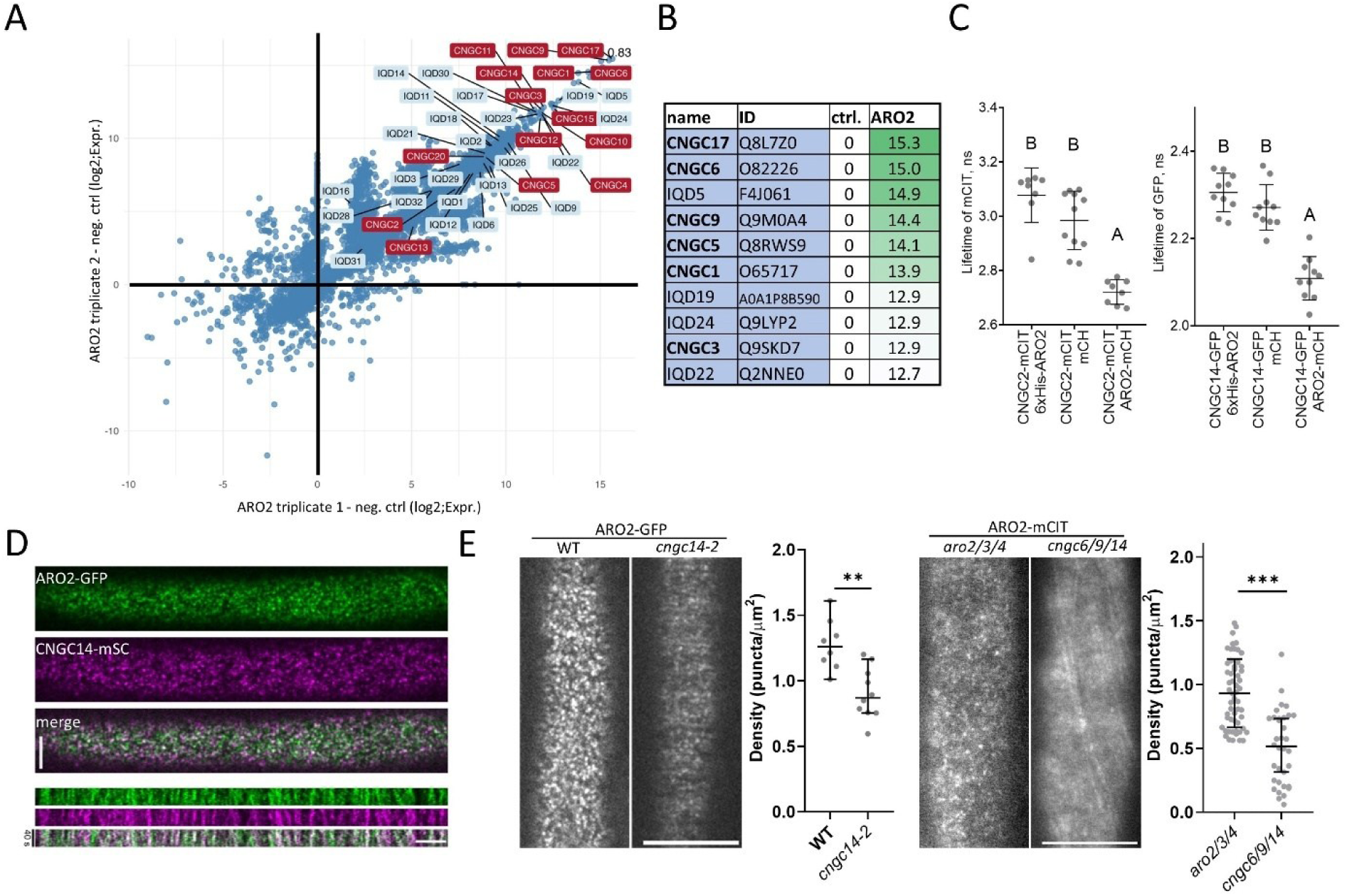
AROs interact with all sporophytic CNGCs. **(A)** Scatter plot highlighting of all CNGCs found as significant targets in ARO2 proxiome. Comparison of two triplicates and negative control. **(B)** Top ten most enriched proteins in ARO2 proxiome compared to neg. Control. Values represent mean binary logarithms of protein intensity and are average of triplicate. **(C)** FRET-FLIM performed in *N.benthamiana* shows positive interaction results for both CNGC14-ARO2 and CNGC2-ARO2 pairs. Letters depict significantly different datasets with p-value <0.0001 (one-way ANOVA). Each measurement represents an individual cell (n=10) **(D)** ARO2 partly co-localizes with CNGC14 in root epidermal cells. Bottom – line intensity profile over 40 seconds of imaging. Bar=5µm **(E)** ARO2-GFP gradually loses its membrane puncta with loss of CNGCs. Left – cortical section using SD confocal microscopy. WT was outcrossed from the *cngc14-2* heterozygote. Right – *aro2/3/4* mutant complemented by ARO2-mCIT compared to ARO2-mCIT in *cngc6/9/14*.

Further, we performed a reciprocal proximity labeling experiment using CNGC2-tID. The top three differentially biotinylated proteins were ARO4, CNGC4 and ARO3. CNGC4 is known to form heterotetramers with CNGC2^15^, confirming the specificity of the experiment (Fig. S1C).

We then selected CNGC2 and CNGC14 as the representative CNGCs, the first being expressed throughout the plant body and involved in response to pathogens or damage^11,27^, the latter as a root-specific CNGC shown to mediate response to the phytohormone auxin^3^. First, we analyzed their interaction with ARO2 using the *Nicotiana benthamiana* transient expression system coupled with FRET-FLIM imaging. In both CNGC2 and CNGC14, the coexpression with ARO2-mCherry (mCH) caused a significant reduction of the CNGC-mCitrine (mCIT) lifetime (Fig. 1C), indicating a direct interaction between the proteins. In root epidermal cells of stably transformed Arabidopsis plants, ARO2-GFP and ARO3-GFP localized to punctae decorating the plasma membrane. CNGC14-mSCARLET (mSC) partly co-localized with ARO2-GFP and with ARO3-GFP punctae, (Fig. 1D, Fig. S1D, Supplementary Movie 1), indicating that these represent plasma-membrane-localized channel complexes. The CNGC14-mSC decorated a subset of ARO2-GFP puncta, which is expected given that AROs interact with multiple CNGCs.

Further, we analyzed the dependence of ARO2 plasma membrane localization on CNGCs. In the *cngc14-2* mutant, the number of ARO2-GFP punctae was dramatically reduced, and the punctae were nearly absent in the *cngc6/9/14* triple mutant expressing ARO2-mCIT (Fig. 1E). Interestingly, the remaining punctae often formed lines resembling microtubules (Fig. 1E).

Given the above, knowing that all AROs are functionally conserved and ancestral, we concluded that AROs interact with all sporophytic CNGCs in plants. Thereby, CNGCs appear to recruit AROs to co-localize within their punctate pattern in the plasma membrane. Gametophyte-specific ARO1 can complement *aro2/3/4* and vice versa^25^ when expressed under control of the ARO2 promoter. Thus, the conclusion can be extrapolated to the remaining 5 gametophytic CNGCs and to CNGCs across other land plants.

### *aro2/3/4* plants phenocopy higher order *cngc* mutants and lack rapid auxin responses

The most notable phenotype of the *aro2/3/4* triple mutant is the bursting of root hairs immediately after bulging, a phenomenon we previously described^25^. In addition, *aro2/3/4* mutant roots exhibit enhanced waving when grown on 45° tilted plates (Fig. 2A). This phenotype closely resembles that of the *cngc14* single and the *cngc6/9/14* triple mutant (Fig. 2A), the latter also displaying the root hair bursting phenotype^6^ (Fig. 3A).

**Figure 2.**
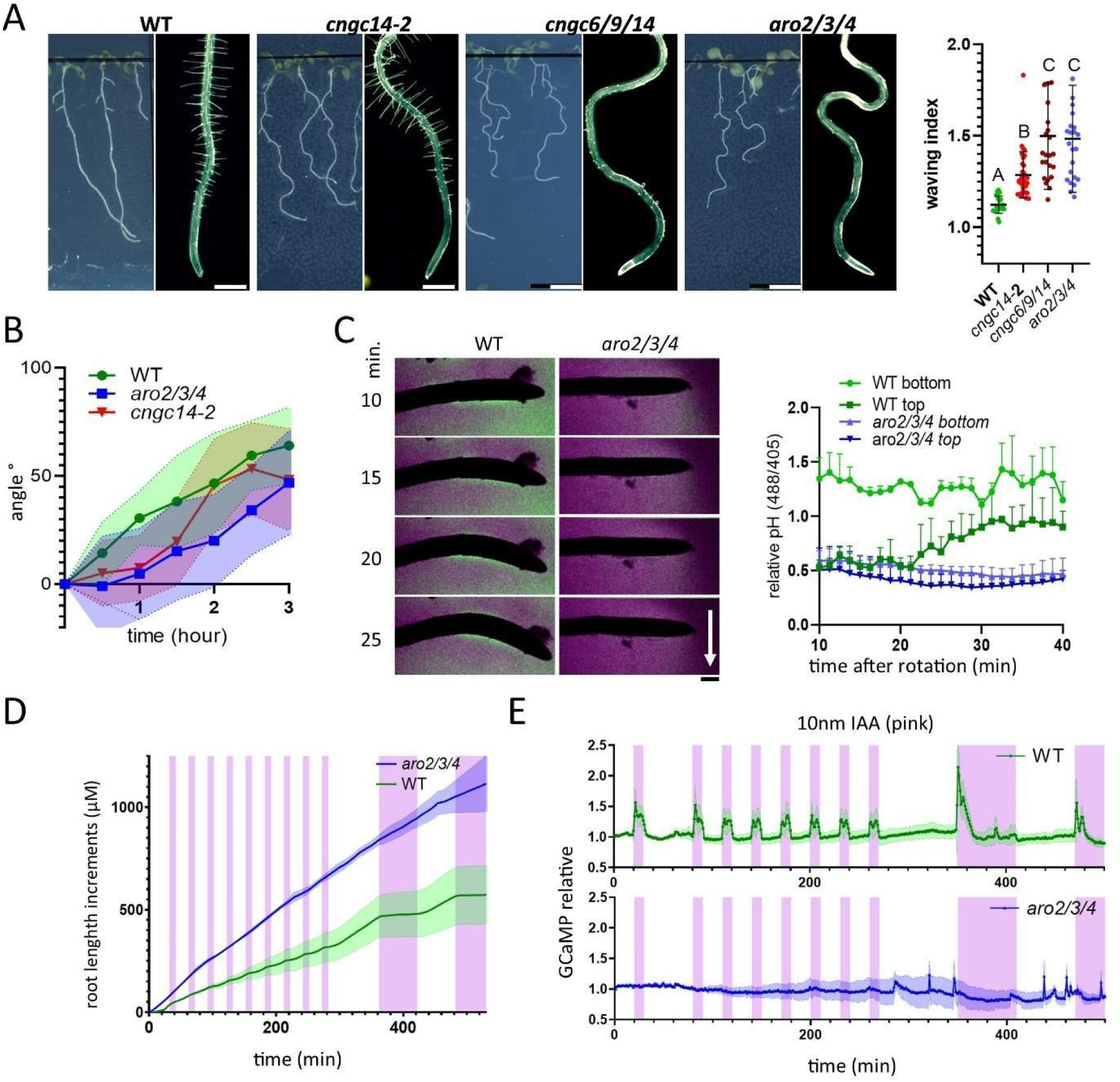
*aro2/3/4* mutants phenocopy higher order *cngc* mutants and lack rapid auxin signaling. **(A)** Representative images depict root waving phenotype on tilted plates (left panel) and root hair growth phenotype (right panel) of *aro2/3/4* in comparison with *cngc14-2* and *cngc6/9/14* triple mutants. 7-day-old seedlings. Bars=500µm. Right-Quantification of the root waving shown as the waving index. Each measurement (n>20) represents an individual root grown on a 45°angle. Error bars = SD. **(B)** Root gravitropic bending of *aro2/3/4* and *cngc14-2* mutants. 14 plants per genotype were evaluated. Similar results were obtained >3 times. Colored areas = SD. **(C)** Apoplastic pH profile of the *aro2/3/4* mutants and WT during the course of the root gravitropic bending, where green halo represents alkaline shift, magenta halo represents acidic shift, and gravitropic vector is depicted by an arrow. Whole movie - Supplementary Movie 3. Bar=50µm. Right - quantification of the pH shifts from (C) on 3 roots per genotype. Two similar replicas were generated. Error bars = SD. **(D)** Kinetics of the root growth inhibition upon auxin treatment. Magenta stripes represent time windows with 10nM IAA. Growth increments were averaged from 2 plants; similar experiments were obtained >3 times. Colored areas = SD **(E)** 10nM IAA treatment fails to induce Ca^2+^ transients in *aro2/3/4* mutants. Pink stripes represent the time windows with the treatment. Average of 3 roots. Two more similar replicas were generated. GCaMP intensity was normalized as follows: (GCaMP^ex488^/GCaMP^ex405^)/baseline (avg of t1-t30). Colored areas = SD. See Supplementary Movie 4.

**Figure 3.**
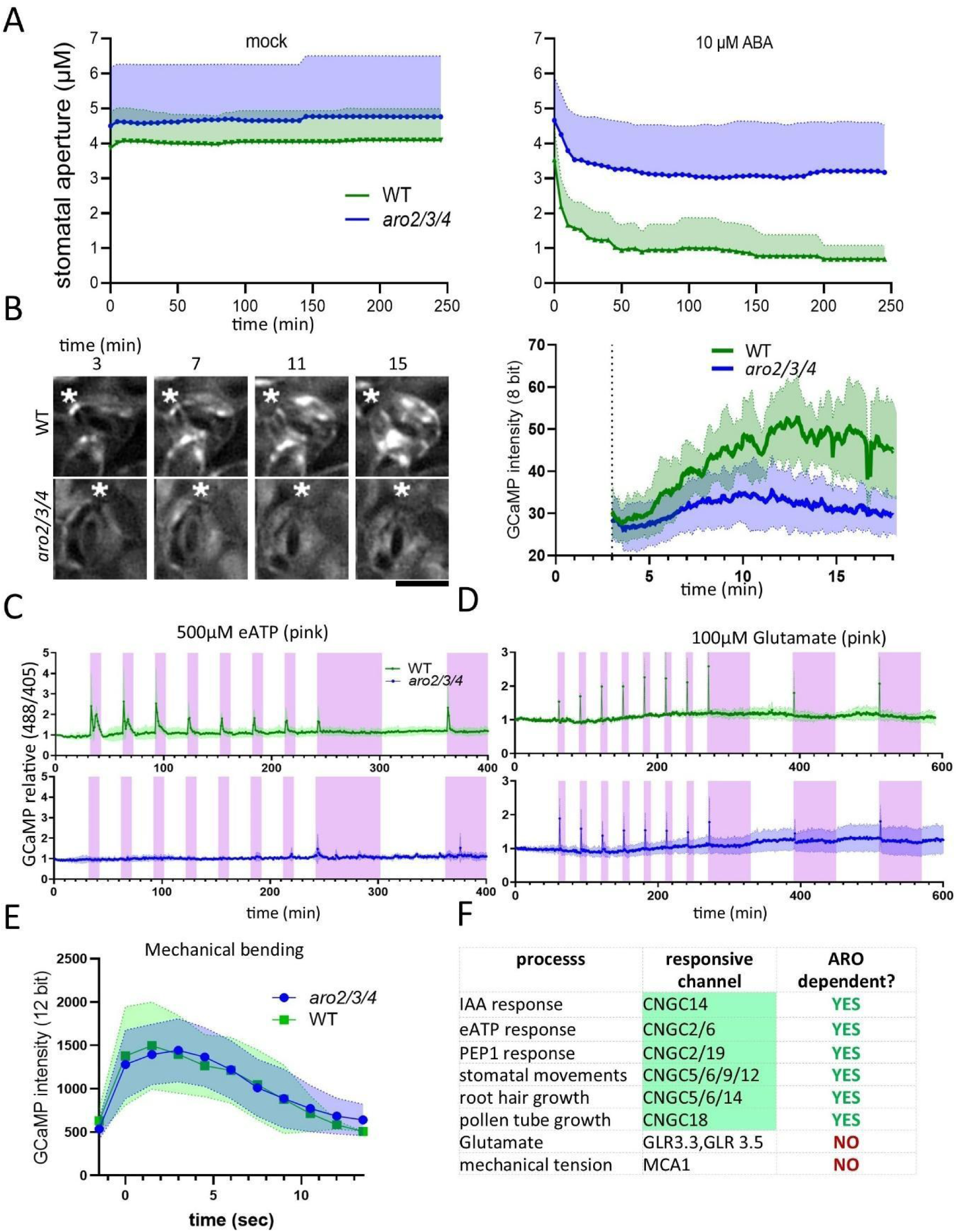
*aro2/3/4* mutants fail to generate CNGC dependent Ca^2+^ transients. **(A)** Quantification of the stomatal aperture after 10µM ABA treatment. 10 stomata per genotype were measured. **(B)** Left- Representative images of WT and *aro2/3/4* undergoing drought stress. Asterisk depicts junction of dorsal guard cell contact sites. Scalebar =10µm. Right – Quantification of the guard cell GCAMP intensity upon drought stress. 16 guard cells were imaged. The experiment was repeated 3 times. See supplementary Movie 5. **(C)** 500µM eATP treatment fails to generate Ca^2+^ transients in *aro2/3/4* mutants. 4 plants per treatment were measured. The experiment was repeated 2 times. **(D)** 100µM glutamate treatment induces comparable Ca^2+^ transients in both *aro2/3/4* and WT. Pink stripes represent the time windows with the treatment. 3 plants per genotype were evaluated. Similar results were observed 3 times. **(E)** Mechanical bending induces Ca^2+^ transient on the convex side of the root in both *aro2/3/4* and WT plants. 6 plants were measured. Experiment was repeated 2 times. **(F)** Summary of the multiple responses and phenotypes of *aro2/3/4* mutants along with responsible Ca^2+^ channels. Colored areas = SD. Pink stripes represent the time windows with the treatment.

CNGC14 mediates the ultra-rapid Ca^2+^ influx in response to auxin^3^, and the *cngc14* mutants lack the auxin-induced Ca^2+^ transient, apoplast alkalinization and display a delay in gravitropic root response^3^. Due to the lack of ARO3 expression in the elongation zone^25^, we detected a comparable root gravitropic defect of *aro2/4* and *aro2/3/4* mutants (Fig. 2B, S2A) that was pronounced in the early phase of the response (Fig. S2B; Supplementary Movie 2). The roots of *aro2/3/4* failed to alkalinize the apoplast at the lower root side (Fig. 2C, Supplementary Movie 3), and lacked the auxin-induced membrane depolarization (Fig. S2C). Importantly, the formation of auxin gradients in the mutant was comparable to the WT control (Fig. S2D). These results indicate a defect in auxin-induced Ca^2+^ transients. Therefore, we introduced the GCaMP^29^ calcium sensor into the *aro2/3/4* mutant, and analyzed its physiological responses in a high spatio-temporal resolution using a microfluidic setup^2^. This revealed that *aro2/3/4* mutants lack auxin-induced rapid root growth inhibition entirely (Fig. 2D), as well as auxin induced Ca^2+^ transients (Fig. 2E, Supplementary Movie 4).

Taken together, the waving and gravitropic bending phenotypes in *aro2/3/4* can be attributed to the absence of Ca^2+^ transients mediated by CNGC14, while the lack of root hair development may reflect a combined effect of CNGC6, 9, and 14.

### AROs are essential for CNGC dependent signaling

To verify the functionality of other CNGCs in the *aro2/3/4* mutant, we examined additional CNGC-dependent processes. First, we focused on stomatal movement as CNGC-mediated Ca^2+^ transients are essential for stomatal closure^4,5^. Indeed, the *aro2/3/4* triple mutant exhibited a severe defect in stomatal closure (Fig. 3A), phenocopying the *cngc5/6/9/12* quadruple mutant^4^. To investigate whether the stomatal defects in *aro2/3/4* are associated with disrupted Ca^2+^ signaling, we subjected 7-day-old cotyledons to drought stress. In WT plants, drought stress induced Ca^2+^ spiking in stomata, whereas these transients were significantly reduced in *aro2/3/4* mutants (Fig. 3B; Supplementary Movie 5).

Second, we analyzed the response to eATP which triggers CNGC2- or CNGC6-dependent Ca^2+^ transients^11,12^. The treatment with eATP failed to elicit a detectable response in *aro2/3/4* mutant roots (Fig. 3C). Similarly, the AtPEP1 peptide, previously identified as an activator of the CNGC2, did not induce Ca^2+^ transients in *aro2/3/4* mutants (Fig. S3A).

Another critical argument for a functional link was demonstrated by overexpressing CNGCs. Expression of CNGC14 under its native promoter in WT plants resulted in slightly, but significantly, shorter roots compared to WT controls (Fig. S3B). However, this effect was not observed in *aro2/3/4* mutants expressing comparable levels of CNGC14, which retained all mutant phenotypes. Similarly, expression of CNGC2 driven by the constitutive ubiquitin-10 (UBQ) promoter showed similar localization in both the *aro2/3/4* and WT backgrounds (Fig. S3C), but caused dwarfism in WT plants while the *aro2/3/4* mutants maintained their original size (Fig. S3D).

In contrast, *aro2/3/4* mutants retained Ca^2+^ transients mediated by other Ca^2+^ channels such as GLR3.3 and GLR3.6, which are activated by glutamate^29^ (Fig. 3D), as well as MCA1-dependent Ca^2+^ transients elicited by plasma membrane (PM) stretching^30^ (Fig. 3E), confirming the specific CNGCs defect in *aro2/3/4* mutants.

In summary, all tested CNGC-dependent Ca^2+^ signaling processes were abolished in *aro2/3/4* mutants, while CNGC-independent Ca^2+^ transients remained intact (Fig. 3F). These confined, stimulus-specific Ca^2+^ response phenotypes indicate that ARO proteins specifically target CNGC-dependent pathways and may be intimately linked to their molecular functions.

### AROs do not primarily act via CNGCs’ stabilization or phosphorylation

To explore the possible mechanism behind the AROs’ action in CNGC operation, we examined the abundance and localization pattern of CNGCs in cells lacking AROs. Both CNGC14 and CNGC2 localized normally in the *aro2/3/4* mutant (Fig. 3C, S4A). Similar signal intensities and similar amounts of PM puncta were visible in both *aro2/3/4* and WT plants expressing CNGC14-GFP (Fig. S4A-C), without affecting the *aro2/3/4* phenotypes (Fig. S4D,E), thus ruling out the stabilization or trafficking role of AROs. Comparison of the CNGC2 proxiome in *aro2/3/4* and WT backgrounds revealed similar amounts for CNGC4, a known heterodimerization partner^27^, as well as for some other CNGCs (Fig. S4E) and known CNGC activating kinases such as BRI1^1^. Single molecule photobleaching experiments confirmed that CNGC14 homotetramers were present in both *aro2/3/4* and WT (Fig. S4F). We conclude that the *aro2/3/4* phenotypes cannot be explained by differential presence or assembly of the CNGC tetramers.

Phosphorylation has been shown as a molecular switch for CNGC activation (shown for CNGC5,6,9,12^4,5,24^) which might either act independently of or in concert with AROs. We generated both wild type and phospho-mimetic versions of a known phosphorylation site (S26) in CNGC9^5,24^, which were expressed under the native promoter fused with miniTurbo-YFP tag. As expected, phospo-mimetic CNGC9^S26D^ did rescue *cngc6/9/14* root hair phenotype to the higher extent in comparison with the non-mutated form (Fig. S5G), whereas *aro2/3/4* mutants were not complemented by either of the two CNGC9 variants (Fig. S4G). This indicates that CNGC9 cannot reach its open conformation without ARO, regardless of its N-terminal phosphorylation status.

### CNGCs interact with AROs via their IQ domain, forming larger molecular complex

We have shown that AROs physically interact with CNGCs on the PM, and the loss of function phenotypes suggest that AROs are required for activation of the CNGC tetrameric channels. To address how AROs interact with CNGCs, we performed structural modeling of CNGC2, CNGC9 and CNGC14 (Fig. S5) tetramers in complex with ARO2 using AlphaFold3^31^ (Fig. 4 and Fig. S5). The model confidently predicted an octameric complex (Fig. 4A,C), where each CNGC monomer recruits one ARO2 via its IQ domain (Fig. 4B), which is the same region binding calmodulins^32^. AROs share the IQ domain with IQD proteins, which we identified in the ARO2 proxiome (Fig. 1A,B), corroborating that the IQ domain forms the ARO-CNGC interaction interface. Additionally, the model is consistent with our previous observation^17^ that the N-terminal basic residues of ARO2 associate with the PM (Fig. 2C).

**Figure 4.**
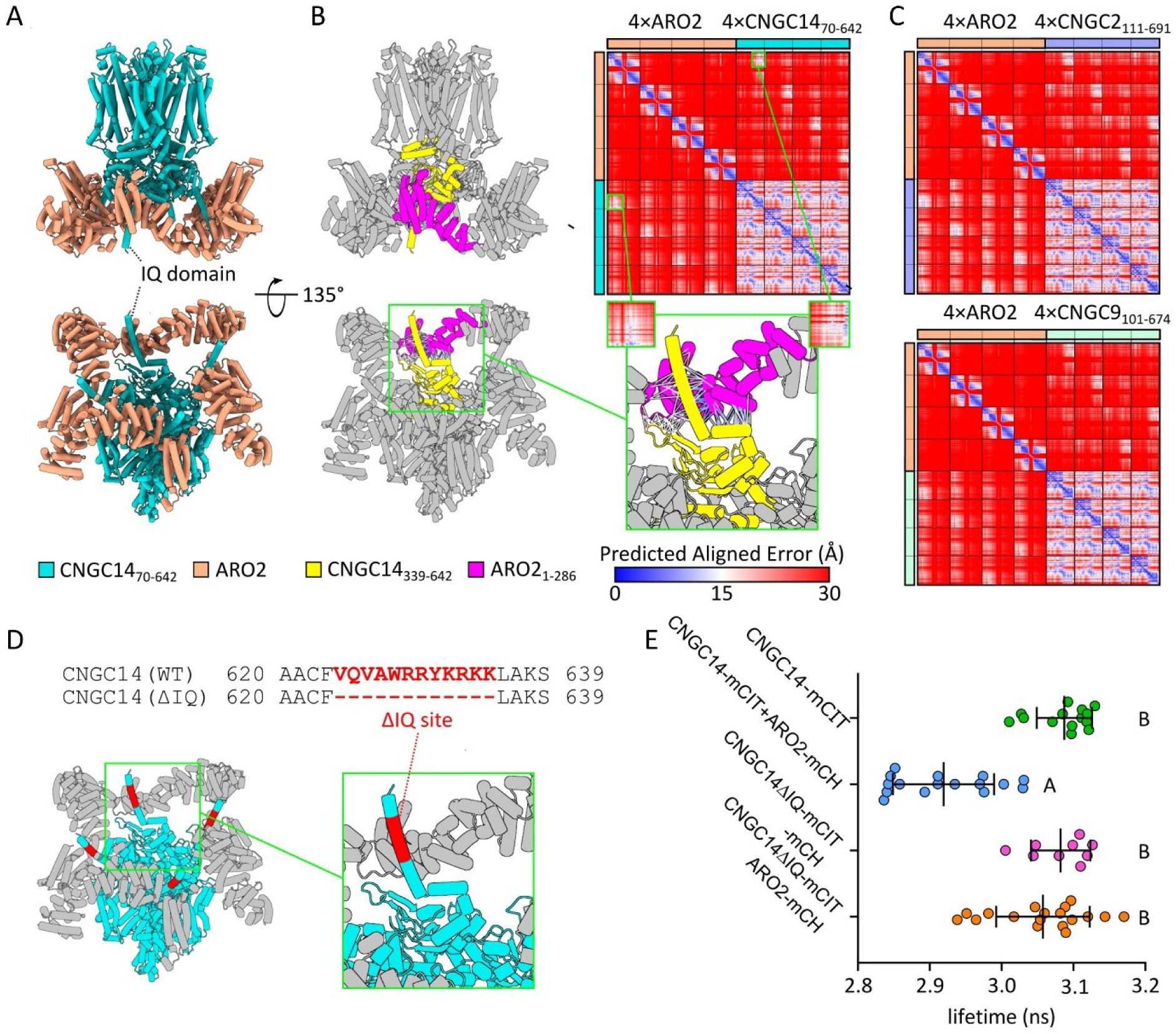
AROs interact with CNGCs via the IQ domain. **(A)** AlphaFold3 prediction of the ARO2-CNGC14 complex. Parts predicted with the low pLDDT score were excluded from the visualization (see Methods and Fig. S5). **(B)** C-terminal part of CNGC14 (IQ domain) and the N-terminus of ARO2 forming the interaction interface are highlighted in yellow and magenta. The Predicted Aligned Error (PAE) plot shows AlphaFold3 confidently predict this interaction. The zoom-in image highlights contacts under 8 Å between these regions. Lines are colored according to the PAE values, with most contacts showing high confidence. **(C)** AlphaFold3 prediction of the ARO2-CNGC2 and ARO2-CNGC9 complex. The PAE plot shows AlphaFold3 confidently predicts these interactions. Corresponding structural models are shown in Fig. S5A, B. **(D)** Depiction of the IQ domain (red) and its deletion (ΔIQ) **(E)** FRET-FLIM measurement of the CNGC-ARO interaction showing IQ domain is essential for the CNGC binding to ARO. Letters depict significantly different datasets with p-value <0.0001 (one-way ANOVA). The experiment was repeated 3 times

To validate the model experimentally, we tested the interaction of ARO2 and CNGC14 with or without IQ domain (CNGC14^ΔIQ^) using FRET-FLIM (Fig. 2D, E). Removal of the IQ domain shifted the fluorescence lifetime to the levels of the negative controls (Fig. 2E), indicating the loss of interaction.

These results confirm that the IQ domain represents the interaction interface between ARO2 and CNGCs, and validate the model, in which AROs form a homo or hetero tetramer surrounding the pore-forming CNGC tetramers.

### AROs activate CNGCs in *Xenopus* oocytes and compete with Calmodulins for binding the IQ domain

To gain insights into the implications of the ARO-CNGC interaction for channel function, we expressed CNGCs in the heterologous *Xenopus* oocyte system. Surprisingly, the co-expression of ARO2 and CNGC14 caused cytotoxicity, not seen in control oocytes, preventing a proper electrophysiological analysis (Fig. S6A). Cytotoxicity is a well-known issue associated with the expression of highly active Ca²⁺ channels in *Xenopus* oocytes^33,34^. These toxic effects upon co-expression with ARO2 were not observed when CNGC14VQ/DA was used, which carries two point mutations in the conserved IQ motif. The oocyte survival rates together with the lack of a stimulatory effect of ARO2 on CNGC14VQ/DA currents (Fig. 5A) suggest that ARO2 activates the CNGC14 channel by binding to the IQ domain. These results corroborate our structural modeling and imply that the IQ domain is not essential for CNGC14 function per se, but indispensable for the interaction with ARO2 and Calmodulin 2 (CaM2). To avoid the CNGC14 toxicity in the oocyte system, we co-expressed ARO2, CNGC14 and the calcium sensor GCaMP in Human Embryonic Kidney (HEK293T) cells. Consistent with our previous observations, the CNGC14-mediated Ca^2+^ entry was dramatically elevated in the presence of ARO2, supporting its role as a channel activator (Fig. 5B).

**Figure 5.**
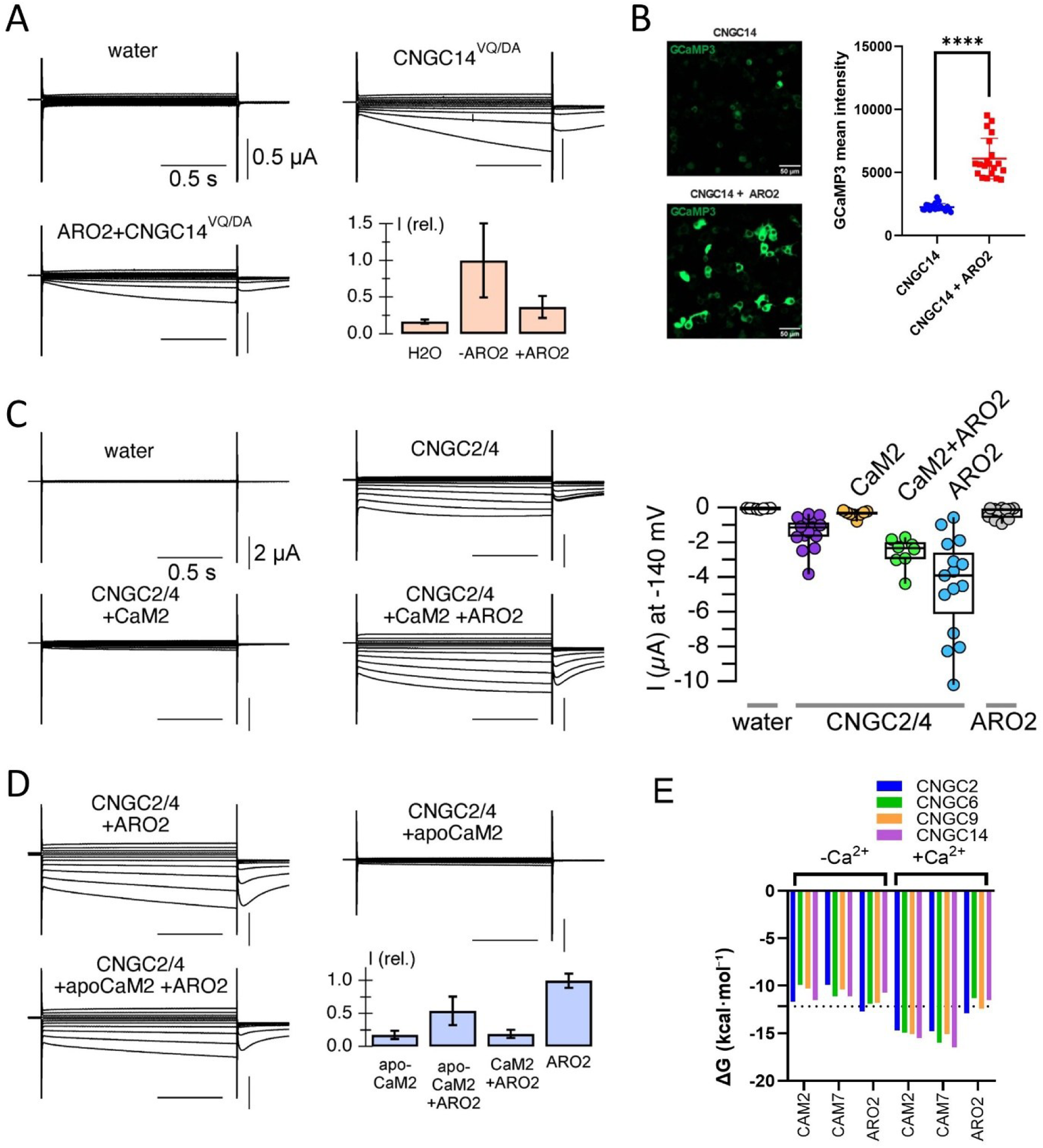
ARO2 activates CNGCs and competes with CaM in a Ca^2+-^dependent manner. **(A)** ARO does not activate the CNGC14VQ/DA mutant. Current traces of oocytes expressing CNGC14VQ/DA in the presence and absence of ARO2. Currents were recorded in 30 mM CaCl_2_+ 500 µM NFA. For CNGC14 currents, see Fig. S6A. **(B)** ARO can activate CNGC14 in HEK cells. Cells were co-transfected by GCaMP and single vector expressing CNGC14 and ARO2 separated by P2A cleavage site. 20 images per each experiment were chosen randomly out of 12 wells. **(C)** ARO2 and CaM2 antagonistically regulate CNGC2/4. *Left:* Current responses of *Xenopus* oocytes injected with water or cRNA of CNGC2 and CNGC4, in the presence and absence of CaM2 and ARO2, as indicated. *Right:* Box plot of current amplitudes at −140 mV of oocytes injected with water (n=8) or ARO2 (n=11) as controls, or co-injected with CNGC2 and CNGC4 together with water (n=18), CaM2 (n=8), CaM2 and ARO2 (n=8), and ARO2 (n=19). Currents in (C) and (D) were recorded in 30 mM BaCl_2_. **(D)** CaM2-dependent interference with ARO-induced activation is reliant on its calcium binding ability. Current responses of oocytes co-injected with CNGC2 and CNGC4 together with apoCaM2 (n=10), or ARO2 (n=10), or ARO2 in the presence of apoCaM2 (n=9) or CaM2 (n=10), respectively. Category plot indicates normalized mean currents at −140 mV (± SEM). **(E)** Changes in Gibbs free energies for complex formation of IQ domain with CaMs. CaM5 and CaM2 share 100% amino acid sequence.

**Figure 6.**
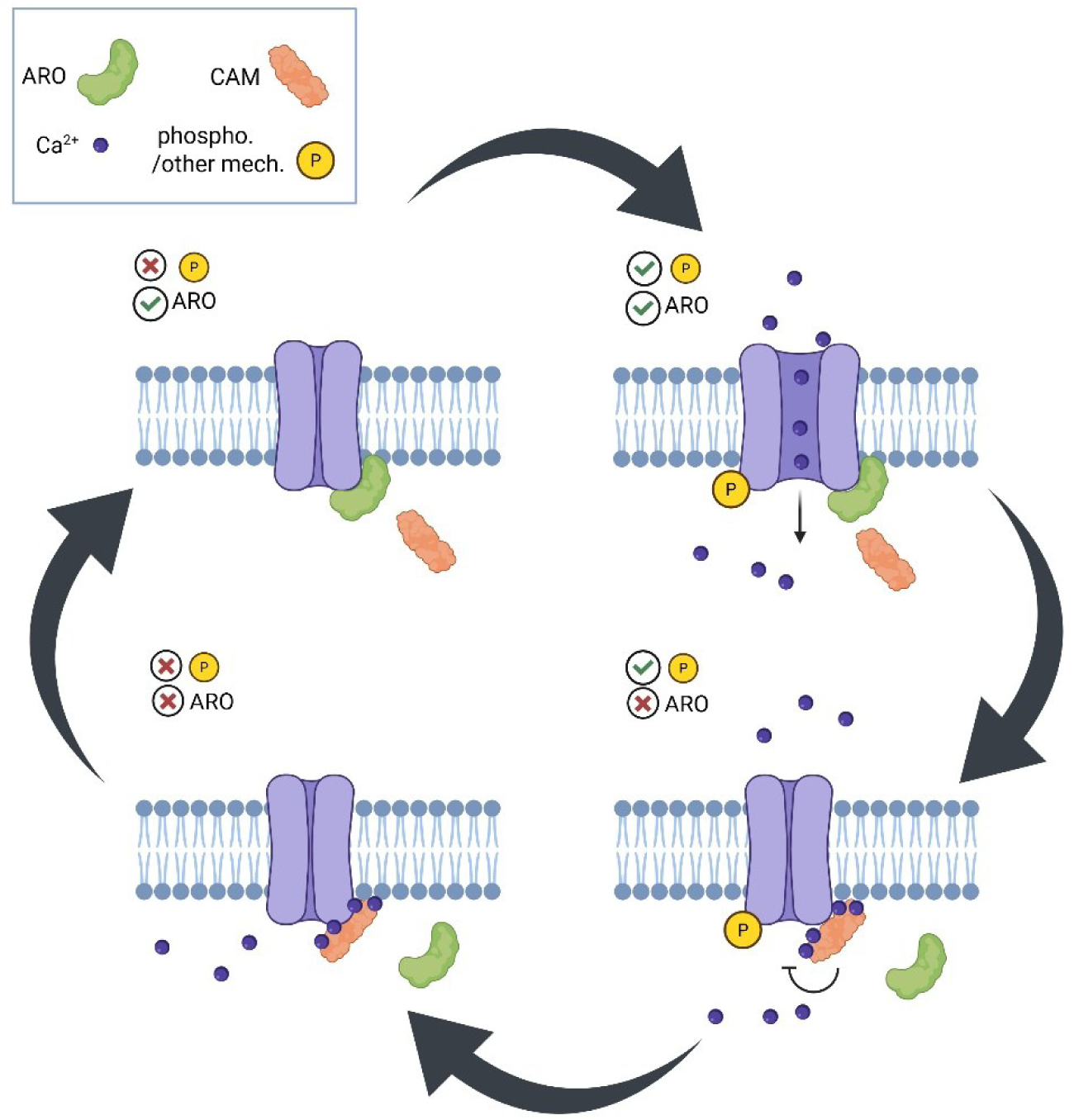
Graphical representation of the ARO dependent CNGC gating.

Further, we used the CNGC2/4 heterotetramer as an additional model for studying the ARO2-CNGC regulation. Two-electrode voltage-clamp (TEVC) recordings confirmed that co-expression of CNGC2 and CNGC4 results in functional channels, which were inhibited by co-expression of CaM2^15,35^ (Fig. 5C). This calmodulin-dependent inhibition was relieved in the presence of ARO2, supporting antagonistic regulation by the two proteins. CNGC activation by ARO2 was also obvious from the increased current amplitudes of CNGC2/4-expressing oocytes upon co-expression of ARO2 in the absence of CaM2 (Fig. 5C,D, Fig. S6B).

Because calmodulin is modulated by binding of Ca^2+^ to its EF-hands, we focused on the role of Ca^2+^ binding in ARO-CaM antagonism. Compared to CaM2, co-expression of an engineered apo-CaM2 (constitutively Ca^2+^-free) could still inhibit the channel function, however, in combination with ARO2, its effect on the current amplitudes was substantially weaker in comparison with WT CAM2 (Fig. 5D). This is in line with our Gibbs free energy calculations, which indicate that the energy required to break the ARO–IQ domain complex is similar or lower to that of the CAM–IQ domain complex in the absence of Ca^2+^. However, the binding of Ca^2+^ enhances CAM’s affinity for the IQ domain (Fig. 5E).

To test if AROs indeed can be displaced by calmodulin *in planta*, we performed FRET-FLIM analysis of ARO2 and CNGC2 as well as ARO2 and CNGC14 in the presence and absence of calmodulin. We selected CAM7, which differs from CAM2 only in a single amino acid, and exhibits the same binding properties (Fig. 5E). To prevent interference of a third fluorophore in the FRET-FLIM experiment, we fused CAM7 to P2A-BFP-NLS, so that the blue fluorophore is cleaved off and transported into the nucleus (Fig. S6C). The addition of CAM7 increased the fluorescence lifetime of CNGC2-mCit coexpressed with ARO2-mCherry. This indicated that the CNGC2-ARO2 complex was destabilized. The destabilization effect was partly reversed by infiltration of 10 mM EGTA about 3 h before imaging (Fig. S6C). Similar trends, but with lower significance were observed in the case of CNGC14 (Fig. S6C)

Taken together, the results support a model, in which CNGCs possess one orthosterical binding site for AROs and calmodulin. When the concentration of Ca^2+^ increases, the affinity of calmodulin towards the channels is increased, favoring the CaM-CNGC complex formation and reverting the activation by AROs.

## Discussion

### AROs – new kids on the CNGC block

This study identifies members of the ARO protein family as central determinants of CNGCs activity in plants. Evidence is drawn from comprehensive proxiome analyses and the absence of canonical CNGC-mediated responses in *aro* mutant backgrounds. Functional assays in *Xenopus* oocytes and HEK cells further established AROs as direct positive regulators of CNGCs. While we have identified additional IQ domain containing proteins (namely IQDs) in the ARO2 proxiome, we did not observe any IQD related phenotype (microtubular or pavement cell lobing)^26,36^ and all the observed *aro2/3/4* phenotypes are explained by loss of CNGCs activity.

Although the present work centers on sporophytic AROs, the data suggest analogous roles for the gametophytic ARO1. ARO1 expression under the ARO2 promoter restores normal function in *aro2/3/4* mutants and vice versa^25^, while loss-of-function *aro1* mutants exhibit pronounced pollen tube defects, including swelling and rupture^37^. Notably, these phenotypes surpass the severity observed in a pollen specific *cngc18* mutant^7^, suggesting that AROs are required for the activity of additional CNGCs in the pollen tube.

### Implications for CNGCs functions in plants

Analysis of *aro2/3/4* mutants brings further implications for the role of CNGC2 in the immune system. The *cngc2* is a dwarf mutant which shows a lesion mimic phenotype^38,39^ (plant equivalent of autoimmune disease). In contrast, *aro2/3/4* mutants have WT size, if cultivated in optimal conditions. In *aro2/3/4* mutants, CNGC2 is still present, although the channel is not functional, demonstrating that CNGC2 presence, not functionality, is important for *cngc2* lesion mimic and dwarf phenotypes, thus CNGC2 integrity is likely guarded by plants immune system. Indeed, we found key regulators of the effector triggered immunity in the CNGC2 proxiome. Further studying *aro2/3/4* mutants may bring a clear distinction between immunity-related and signaling roles of CNGC2 and maybe also other CNGCs.

Previously, we demonstrated that AROs are essential for the ROP GTPase cycle^25^. However, our current findings reveal that most *aro2/3/4* mutant phenotypes can, in fact, be attributed to the absence of CNGC activity. This novel discovery suggests that the regulation of the ROP cycle by AROs is likely indirect and mediated through Ca²⁺ signaling. If this hypothesis holds true, ROP2, which we previously found to be mislocalized in the *aro2/3/4* mutant, should also be mislocalized in the *cngc6/9/14* mutant. This is indeed what we observed (Fig. S6D), supporting this notion. Therefore, CNGCs and Ca²⁺ signaling as such play an important role in the maintenance of plant cell polarity.

### CNGCs– Doors With Two Locks

The propagation of calcium signals in plant cells is a tightly regulated process, as excessive signaling culminates in cell death^40^. This is why channels such as CNGCs need to be tightly regulated. Here, we propose a “two lock on one door” model - where at least two conditions must be met to open the channel. Various stimuli trigger CNGC activation. CNGC9 is N-terminally phosphorylated in response to OST1 in stomata in response to abscisic acid or CPK1 in the root hair. CNGC14 is activated in an auxin- and AFB1-dependent manner^3,41^, by an unknown mechanism. Nevertheless, these inputs require the second, independent condition - the presence of AROs on the C-terminally located IQ domain.

The mechanism of ARO action involves competition with calmodulins. Calmodulins compete with AROs for CNGC binding; with increasing intracellular Ca^2+^, calmodulins will outcompete AROs and close the channel. The question remains whether AROs are primarily positive regulators of CNGC channels, or rather negative regulators of inhibitory calmodulins. Our electrophysiological and HEK cell results suggest AROs as primarily positive regulators, therefore the lack of CNGC functionality in *aro2/3/4* cannot be attributed to constitutive CAM binding.

Besides calmodulins, additional proteins likely modulate CNGC activity, and AROs themselves are a subject of regulation. When we compared CNGC2 proxiome in WT and *aro2/3/4* mutant, not many additional proteins were absent from the complex in *aro2/3/4* suggesting that AROs do not scaffold additional proteins to form the channel complex. For example, BRI1, a known activator of CNGC2, was highly enriched in CNGC2 proxiomes regardless of ARO presence. This is supported by the experiments in heterologous systems, where AROs were sufficient to activate the channels. Very likely, this situation did not represent a fully open channel, which would result in cell death such as with CNGC14-ARO2 combination, but a channel that is more prone to be opened due to lack of one regulatory mechanism. With similar logic, phosphomimetic CNGC9 does not kill Arabidopsis, because the channel activity is still controlled by the calmodulin-ARO module.

### CNGCs - Beyond the Core Channel

In plants so far, no channels were shown to require additional auxiliary subunits for functioning. Apart from phosphorylation by kinases^5,24^ and modulation by calmodulins^32^ or coupled channels^42^, AROs are the first such auxiliary protein. In animals, such protein complexes are more common, especially in the context of neurobiology, where for example glutamate receptors of the AMPA-type (AMPARs) displays architecture of a defined core and variable periphery, where core contains auxiliary subunits TARPs^43^. Another example would be voltage gated sodium channels which contain regulatory beta subunit^44^. Since AROs are required for the channel function in plants, they also belong to the channel core. Discovery of such an auxiliary, plant-specific CNGC subunit represents a key milestone in plant cell signaling and highlights an evolutionary diversification between plants and animals in controlling key signaling processes.

## Materials and Methods

### Plant growth and conditions

All *Arabidopsis* mutants and transgenic lines which were used in this project are in the Columbia-0 (Col-0) background. *aro2/3/4* and *aro2/4* represent T-DNA insertional lines described previously^17^. *cngc14-1* was described in 19 and crossed with ARO2-GFP from 17. *cngc14-*2 were obtained from^3^ and *cngc6/9/14* were obtained from^6^. CNGC14-mScarlet-I was introduced into ARO2-GFP and ARO3-GFP (17) by transformation. *aro2/3/4* plants were crossed into the plants expressing GCAMP3 obtained from^24^. WT expressing GCAMP3 was obtained as a sister plant from the cross with *aro2/3/4* displaying similar signal intensity. *aro2/4* was crossed into R2D2 background^36^ and verified by genotyping as previously^17^. Seeds were sterilized overnight using chlorine gas and sown on solid agarose medium consisting of half-strength Murashige and Skoog medium with or without B5 vitamins (Duchefa M0231) (with 1% sucrose (AM+) and 0.8% phyto agar, adjusted to pH 5.9. The sown seeds were stratified at 4 °C for 2 days, then grown vertically at 21 °C under a 16-hour light/8-hour dark cycle.

### Plant treatments

Treatments with ATP (Thermo Scientific R0441), IAA (Duchefa I0901), glutamate (Sigma; PHR2634-1G) were performed in the microfluidic setup with an equivalent amount of solvent added to the mock control. PEP1 peptide (ATKVKAKQRGKEKVSSGRPGQHN; Chinese Peptide Co., LTD.) was used for a direct treatment without microfluidics.

### Gravitropic and mechanical root bending assays

Gravitropic bending experiments were conducted using a vertically oriented scanner setup. Four-day-old seedlings were transferred to a fresh solid agarose medium plate, with roots aligned to maintain a straight orientation. Images were captured at 30-minute intervals and registered using the Fiji StackReg plugin^37^. Root tracking and angle increment calculations were performed manually in Microsoft Excel. Mechanical bending was performed by positioning the root on a microscopy slide in a droplet of medium. One cover slip was placed alongside the root, while another was positioned over the sample, allowing gentle pressure to be applied by moving the first cover slip towards the root during imaging. This setup was mounted onto the microscope stage and allowed to rest for 5 minutes for recovery. During imaging, the inner cover slip was gradually pressed in until the root exhibited visible bending (15-30°).

### ABA treatment

ABA-induced stomatal closing was performed as follows. Lower epidermal cells were peeled from 7th or 8th rosette leaf of 4-week-old *Arabidopsis*. The peeled epidermal cells were placed in buffer (30 mM KCl, 10 mM MES, 50 µM CaCl2, pH 6.15) one night in advance and kept under plant growth condition (150 µmol/s2/s, 16 hours light /8 hours dark) until the following day. After 3 hours into the circadian lighting cycle, the stomata were allowed to open. 10 µM ABA was added in the buffer before starting the imaging. Live stomatal images were captured using a Nikon Ti2E microscope with 40 x objective at 5 min/frame for 4 hours. The stomatal width was measured by Fiji software, with 10 stomata measured per each treatment.

### Drought stress

Young *Arabidopsis* seedlings grown on the 1/2MS plates were placed on the stage of Plants Axioplan 2 Zeiss Fluorescence Microscope. The lid was removed, reducing relative humidity from 100% to the room humidity (∼40%). Imaging was done for 30 minutes during which plants started to exhibit stomatal Ca^2+^ transients. Whole leaves were imaged with ∼480nm excitation and 500-550nm emission filter.

### Molecular cloning and plant transformation

Most of the plasmids generated in this study were generated using the Greengate system ^38^. Greengate building blocks were either obtained (source indicated in the primer list) or cloned using the Gibson assembly (NEBuilder® HiFi DNA Assembly, NEB #E5520) of the backbone fragment and the insert fragment. If BSAI sites were present, multiple fragments were used, and compatible cohesive ends were used for mutagenesis.

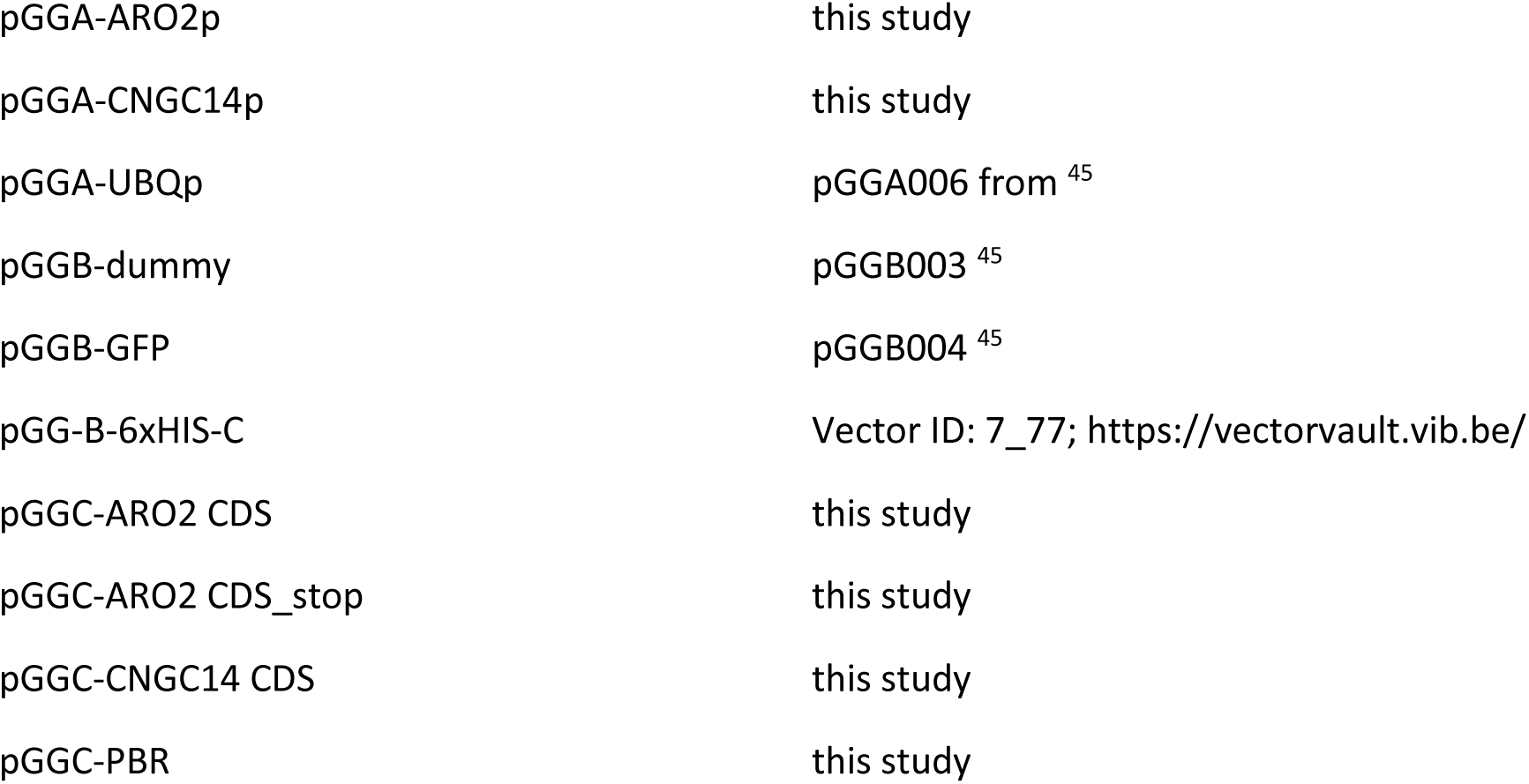
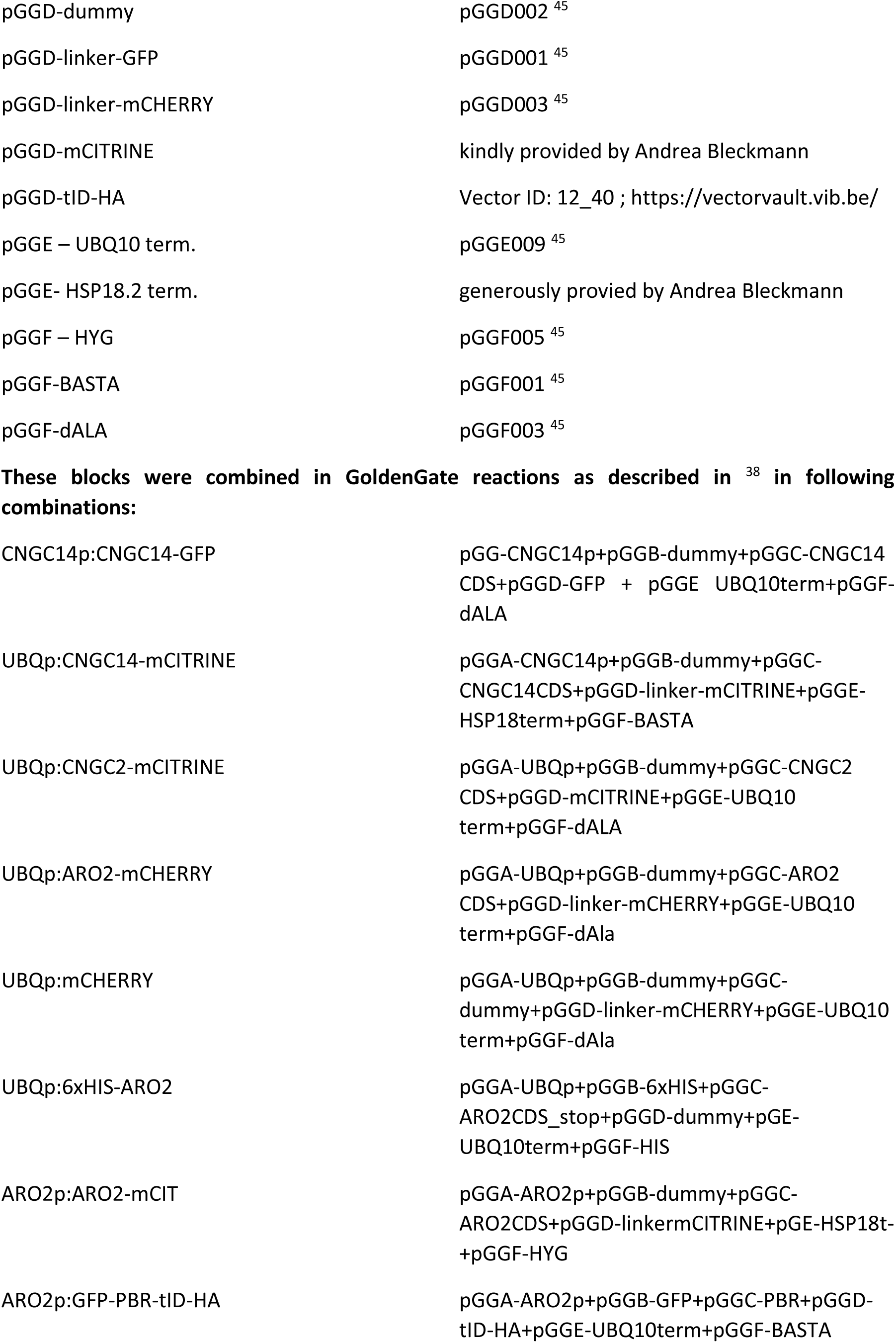
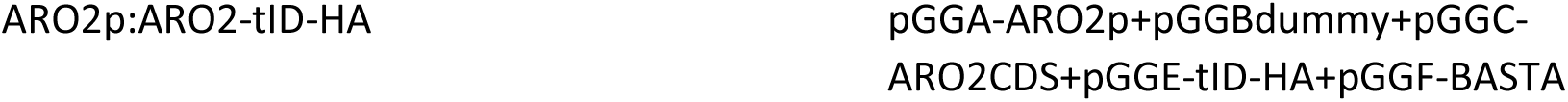

pCNGC14:CNGC14-mScarlet-I binary vector with BaR cassette for plants was assembled using the Golden Braid 2.0^46^ system as described in^47^ for the GFP variant. The CAGp:CNGC14-P2A-ARO2-mCHERRY plasmid was constructed using a combination of Gibson assembly and SalI restriction-based cloning of the vector fragment pBR322. For x. oocyte expression, CNGC2, CNGC4, and CNGC14 were subcloned into modified pGEMHE-GW vectors^48^ via Gateway® technology, carrying tags for the Venus-N (VYNE) or Venus-C (VYCE) part of the split Venus system (Gehl et al. 2009). A tag-free Version of CaM2 was cloned into the pGEMHE-GW. To abolish Ca^2+^-binding ability, a Glu to Ala point mutation was introduced into each EF-hand of CaM2 via PCR, resulting in CaM2E1-4A. CNGC14VQ/DA was produced via mutagenesis PCR. ARO2 and CaM2E1-4A (apoCaM2) were introduced into a pSEVA based oocyte expression vector, using the Golden Gate method^49^.

### Imaging and microfluidics

For the ARO-CNGC colocalization, vertical spinning disc microscope (vertical SD) Zeiss Axio Observer 7 coupled to a Yokogawa CSU-W1-T2 spinning disk unit with 50 µm pinholes and equipped with a VS-HOM1000 excitation light homogenizer (Visitron Systems), operated by the VisiView software (Visitron System) and with alpha Plan-Apochromat 100x/1.46, was used. The same microscope equipped with 5x/0.16 objective was used for observation of the root gravitropic bending (Supplementary Movie 2) and for both the DII-Venus imaging and membrane depolarization assays, with Plan-Apochromat 20x/0.8 M27 obj.

For the Ca^2+^ measurements, vertical stage Zeiss LSM800 (vertical LSM) was used with air objective Plan Apochromat 10x/0.45 M27. GCaMP measurements were normalized by dividing 488nm-induced emission by 405nm induced emission, both between 500-550nm.

Microfluidic imaging was performed as introduced in^2^. This was used either for the membrane potential measurements^39^ with vertical SD and Plan-Apochromat 20x/0.8 M27 with or for the treatments with individual elicitors and vertical LSM setup. For the mechanical bending experiments, Olympus X1000 confocal microscope with 10x dry objective was used. For the stomatal imaging, Axioplan 2 Zeiss Fluorescence Microscope was used.

### FLIM imaging of *Nicotiana benthamiana* leaves

Transiently transformed *N. benthamiana* leaves with A. tumefaciens GV310140 (2 days post-transformation) were mounted on the glass slide and imaged, using the confocal microscope FV3000 (Olympus) equipped with the 40x UApo N water immersion objective lens NA 1.15 (Olympus), MultiHarp 150 Time-Correlated Single Photon Counting device, pulsed 488 nm laser, and two Hybrid Photomultiplier detectors (all Picoquant). The donor fluorophore was excited with the 488 nm laser, working in a pulsed mode with the 40 MHz frequency, emitted fluorescence was filtered using the FF02-525/40-25 emission filter (Semrock), and the signal was detected by a HyD detector, working in the Counting mode. Data analysis was done using Picoquant SymphoTime64 software package. Membrane regions were manually selected in the images, and lifetime (τ) values were calculated from the decay curves using the exponential tailfit with the fixed time window for analysis. The acceptable goodness of fit χ2 values were between 0.9 and 1.2.

### TIRF microscopy and single particle tracking

Total Internal Reflection Fluorescence (TIRF) microscopy was performed on the Elyra 7 microscope (Zeiss) equipped with a 488 nm laser, a 63x Plan-Apochromat objective lens NA 1.46, and a sCMOS camera. Images were acquired every 50 ms (20 frames per second). Individual particles in the images were detected in Fiji ^50^using the TrackMate plugin^51^ with the LoG detector, detection diameter set to 0.324 µm, and a standardized quality threshold. Cell outlines in the TIRF field were manually selected and the density of particles was presented in particles/µm^2^ of the cell surface. Individual bleaching steps were calculated from molecular traces using the Step Finder script in MATLAB ^52^.

### Proximity labelling

Plants expressing proteins tagged with a biotin ligase (HA-TurboID) at the C-terminus, or ARO2p::PBR-GFP-TurboID, were used. The T3 generation was selected based on comparable anti-HA or YFP signal intensity to the negative control. The protocol of^53^ was followed with the following modifications: plants were grown vertically on 1/2 MS medium. Prior to harvest, seedlings were treated at room temperature with 50 μM biotin for 3 hours. Entire seedlings were collected and incubated for 1 hour on a rotary shaker in extraction buffer EB (100 mM Tris-HCl pH 7.5, 2% (w/v) SDS, 8 M urea). Samples were centrifuged twice for 20 min at maximum speed, and the supernatant was filtered through an RC25 0.45 μm syringe filter. Excess biotin was removed using a PD10 desalting column pre-equilibrated with binding buffer BB (100 mM Tris-HCl pH 7.5, 2% (w/v) SDS, 7.5 M urea). Proteins were eluted by adding 3.5 ml of EB and collected into a 5 ml LoBind tube containing equilibrated streptavidin magnetic beads (50 μl per 1 mg fresh weight sample), then incubated overnight on a rotating device. The next day, streptavidin-Sepharose beads were separated using a magnetic stand. Five washing steps with 800 μl of BB were followed by incubation in a high salt buffer (1 M NaCl, 100 mM Tris-HCl pH 7.5) for 30 min. After two washing steps with Milli-Q water and four washing steps with 50 mM Tris pH 8.0, excess liquid was removed and the wet beads were frozen in liquid nitrogen prior to MS-MS analysis.

### Mass spectrometry analysis after proximity labelling

#### Protein Digestion

Solution with beads from imunnoprecipitation was mixed with 1 M TEAB (Triethylammonium bicarbonate) in 20% SDC (sodium deoxycholate), 100 mM TCEP (Tris(2-carboxyethyl)phosphine) and 500 mM CAA (chloroacetamide) and shaked for 30 min at 60°C. After incubation IP beads were digested in 50 mM TEAB at 37 °C with 0.75 µg of trypsin overnight. After digestion, samples were centrifuged and the supernatant was acidified with TFA to 1% final concentration. Supernatant was washed 3times with ethylacetate and the residual ethyacetate was evaporated. Samples were acidified with TFA to 1% final concentration and peptides were desalted using in-house made stage tips packed with C18 disks (Empore) according to^54^.

#### nLC-MS 2 Analysis

Nano Reversed phase columns (Ion Opticks, Aurora Ultimate TS 25×75 C18 UHPLC column) were used for LC/MS analysis. LC analysis is performed on Vanquish NEO (Thermo Scientific). Mobile phase buffer A was composed of water and 0.1% formic acid. Mobile phase B was composed of acetonitrile and 0.1% formic acid. Samples were loaded onto the trap column (C18 PepMap100, 5 μm particle size, 300 μm x 5 mm, Thermo Scientific) in loading buffer composed of water and 0.1% formic acid. Peptides were eluted with Mobile phase B gradient from 2% to 35% B in 19.7 min. Eluting peptide cations were converted to gas-phase ions by electrospray ionization and analyzed on a Thermo Orbitrap Astral (Thermo Scientific) by data independent aproach. Survey scans of peptide precursors from 380 to 980 m/z were performed in orbitrap at 120K resolution (at 200 m/z) with a 5 × 10^6^ ion count target. DIA scans were performed in astral analyzer. AGC target was set to 5 × 10^4^ and maximum injection time to 3 ms. Precursor mass range 380 – 980 m/z was covered by 4 m/z windows. Activation type was set to HCD with 25 % collision energy.

#### Data analysis

All data were analyzed and quantified with the Spectronaut 19 software^55^ using directDIA analysis. Data were searched against *Arabidopsis thaliana* database (downloaded from Uniprot in May 2024, containing 27 448 entries). Enzyme specificity was set as C-terminal to Arg and Lys, also allowing cleavage at proline bonds and a maximum of two missed cleavages. Carbamidomethylation of cysteines was set as fixed modification and N-terminal protein acetylation and methionine oxidation as variable modifications. FDR was set to 1 % for PSM, peptide and protein. Quantification was performed on MS2 level. Precursor PEP cutoff and precursor and protein cutoff was set to 0.01, protein PEP was set to 0.05. Data were exported and data analysis was performed using Perseus 1.6.15.0^56^

#### IP-MS

Lines expressing comparable amounts of ARO2-tID-HA and PBR-GFP-tID-HA were selected. 7-day old seedlings were homogenized using liquid nitrogen and immediately, extraction buffer was added (: 25 mM 2-amino-2-(hydroxymethyl)-1,3-propanediol hydrochloride (Tris–HCl) pH 7.8, 20 mM MgCl_2_, 15 mM EDTA, 75 mM NaCl, 1 mM TCIP, 1 mM NaF, 0.5 mM NaVO_3_, 15 mM β-glycerophosphate, 0.5 mM phenylmethanesulfonyl fluoride (PMSF), protease inhibitor cocktail, 0.1% Tween, 10% CHAPS. Mixture of buffer and powder 2:1 was then mixed for 1 hour at 4°C using a rotatory shaker and then centrifuged at 4°C and 20 min at 20 000x g. To the supernatant, 35ul of antiHA magnetic beads (Pierce Anti-HA Magnetic Beads, Thermo Scientific; 88837) were added and incubated for 2 hours at 4°C while rotating. Then, 3 washes were done with IP buffer (same as extraction buffer, but lacking buffer protease inhibitors, PMSF, NaF, NaVO3, β -glycerophosphate, CHAPS). This was followed by 3 washes with the wash buffer (50 mM NaCl + 20 mM Tris). 5µL of the resulting beads were used for the quality check using SDS-PAGE and total protein visualizaiton. Resulting beads were subjected to the „on bead“ digest by trypisn. All samples were processed using the iST kit (PreOmics GmbH) using the manufacturer’s modified protocol for on-paramagnetic beads digest. Tryptic digestion was stopped after 1 h and cleaned-up samples were vacuum dried. Finally, samples were re-dissolved with 10 min sonication in the iST kit’s LC LOAD buffer.

Raw files were searched in DiaNN version 1.8.1 in library-free mode against an *Arabidopsis thaliana* proteome sourced from UniprotKB. Match-Between-Runs was turned off. Fixed cysteine modification was set to Carbamidomethylation. Acetyl (protein N-term) was set as variable modification. Data was filtered at 1% FDR.

DiaNN’s output was re-processed using in-house R scripts, starting from the main report table. Peptide-to-protein assignments were checked, then Protein Groups were assembled and quantified using an algorithm which: i) computes a mean protein group-level profile across samples using individual, normalized peptidoform profiles (“relative quantitation” step), then ii) following the best-flyer hypothesis, normalizes this profile to the mean intensity level of the most intense peptidoform (“unscaled absolute quantitation” step). Only unique and razor peptidoforms were used. Protein group-level quantitative values were normalized using the Levenberg-Marquardt procedure.

#### Expression in *Xenopus* oocytes and electrophysiology

In vitro transcription and current recordings were performed as described^57^. Oocytes were injected with water or cRNAs in a total volume of 50nL using the Nanoject III (Drummond Scientific Company, Broomall, PA, USA). For CNGC2/4 co-injection experiments, each cRNA in the mixture was adjusted to a final concentration of 12,5 ng for VC-CNGC2, CNGC4-VC, ARO2, CaM2, and apoCaM2. 28 ng CNGC14-VC or CNGC14-VC cRNAs were injected, together with RNase-free water or 11 ng ARO2 or CaM2. Injected oocytes were stored at 17°C in ND96 solution (96 mM NaCl, 2mM KCl, 1mM CaCl_2_, 1 mM MgCl_2_, 5 mM HEPES-NaOH, pH 7.4), supplemented with 25 µg/ml gentamycin. Ion currents were studied by two-electrode voltage-clamp (TEVC) using the Turbo TEC-10CX amplifier (npi electronic GmbH, Tamm, Germany). Current responses were studied at test potentials between +40 and −160 mV. Oocytes were constantly perfused with a bath solution containing 30 mM BaCl_2_, 10mM HEPES-Tris pH7.4. Alternatively, 30 mM CaCl_2_, 2 mM KCl, 10 mM HEPES-Tris pH7.4 was used in the absence or presence of niflumic acid (NFA) to minimize Ca^2+^-activated anion currents. Solutions were adjusted to 220 mosmol kg-1 using D-sorbitol.

#### Transfection of HEK293T cells

HEK293T cells were cultured in DMEM/F12 medium (Thermo Fisher, Cat. No. A4192002) supplemented with 10% fetal bovine serum (FBS; Thermo Fisher, Cat. No. A5670402) and penicillin/streptomycin antibiotics (Thermo Fisher, Cat. No. 15070063). Two days prior to transfection, cells were seeded into 96-well round-bottom plates (Cellvis, Cat. No. P96-0-N) at a density of 2.0·104 cells per well. Transfection was performed using Lipofectamine 2000 reagent (Thermo Fisher, Cat. No. 11668027) according to the manufacturer’s protocol. Per 12 wells, 2.4 µg plasmid DNA and 6 µL Lipofectamine 2000 were used; for co-transfection of two plasmids, 1.2 µg of each plasmid was used. Cells were imaged by confocal microscopy 48 hours post-transfection.

#### Structural modeling

ARO2-CNGC2/9/14 complexes were predicted via AlphaFold3 web server^31^. The Alphafold3 token limit only allowed to model a truncated complex containing full-length ARO2 and residues 70-642 of CNGC14, residues 111-691 of CNGC2 and residues 101-674 of CNGC9 respectively. Additionally, residues 287-389 and 642-651 of ARO2 were removed from the predicted structures (long stretches with pLDDT score below 50) for the visualisation in Fig. 4 and Fig. S5 and further analysis. The truncated version of the ARO2-CNGC14 complex was positioned in the model plant plasma membrane using the PPM 3.0 Web Server (https://opm.phar.umich.edu/ppm_server3_cgopm, ^58^). Furthermore, the C-terminal regions of CNGC2, CNGC6, and CNGC9—including the beginning of the calmodulin-binding domain (CaMBD), the IQ motif, and adjacent residues—were modeled in complex with calmodulin 2 and calmodulin 7 using AlphaFold3. For each complex, two structural variants were generated to represent calmodulin in its Ca²⁺-bound and Ca²⁺-free conformations. The resulting protein–protein complexes were subsequently evaluated for their predicted binding affinities (ΔG) using the PRODIGY web server^59,60^, which estimates binding free energies from structural interface features

#### Primers used in the study

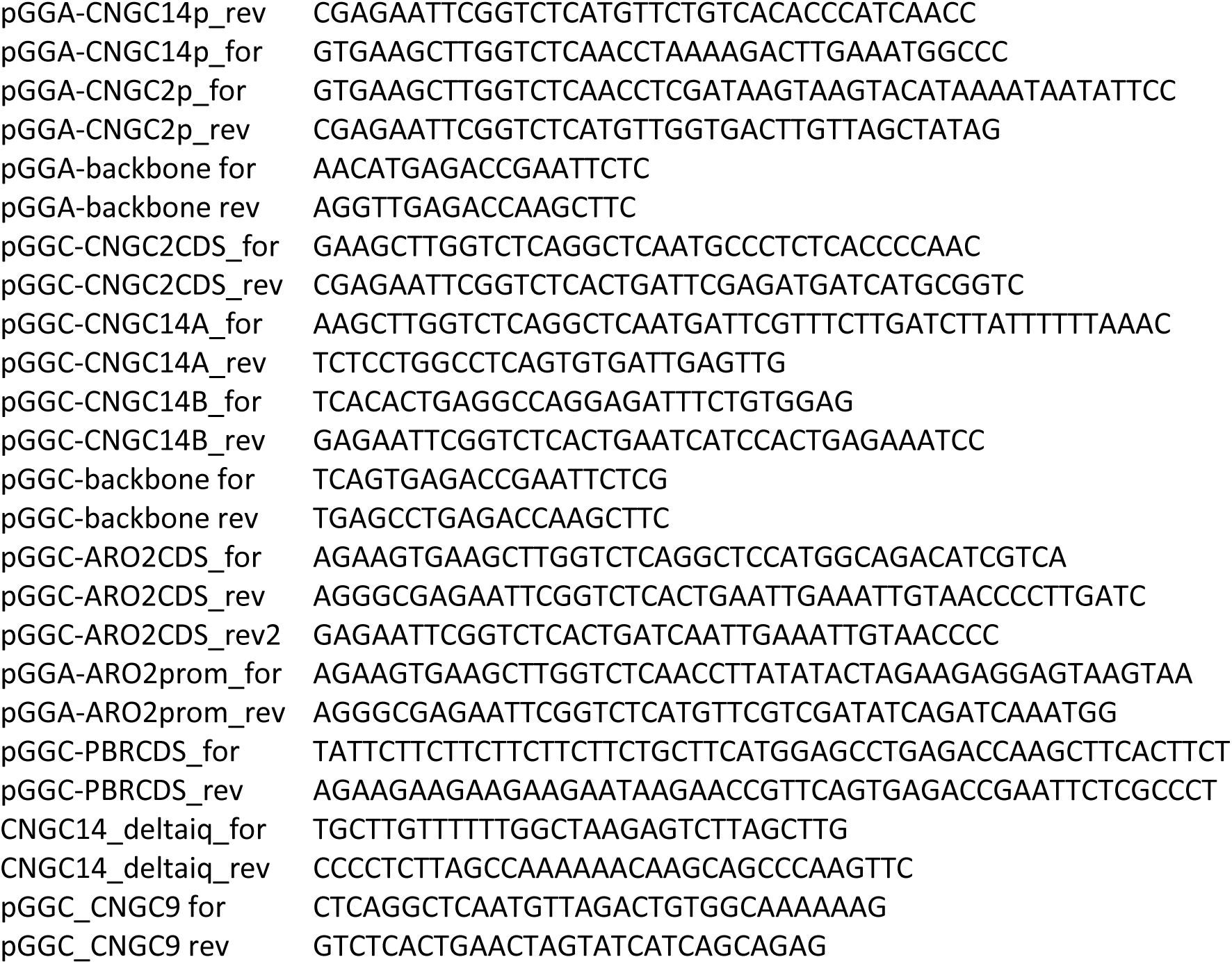
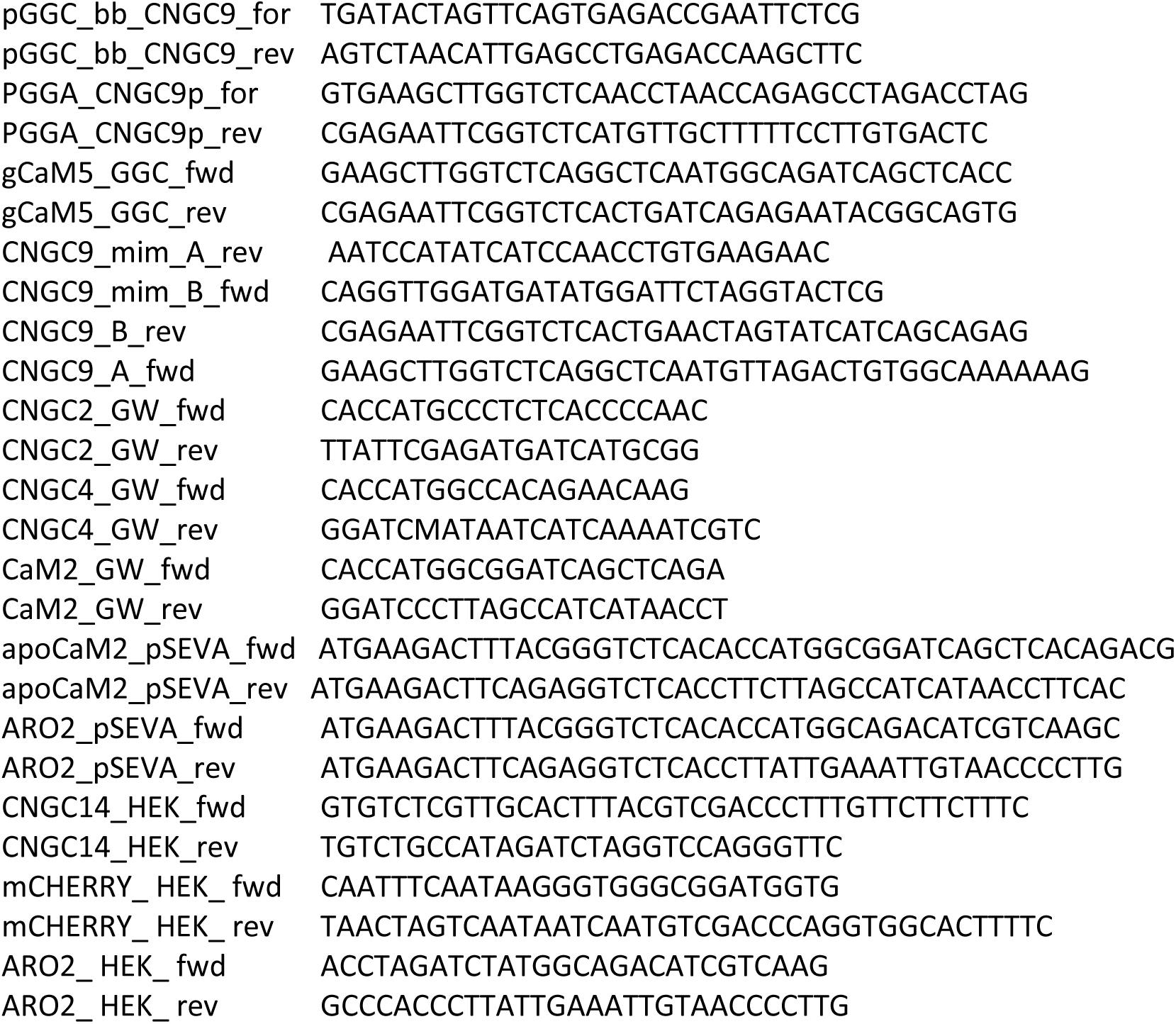

## Acknowledgements

This project was supported by the Czech Science Foundation grant Nr. 25-16449S and by European Union, Horizon Europe, project MOLIPEC, ID 101087030. Computational resources used for structural modeling were provided by the e-INFRA CZ project (ID:90254), supported by the Ministry of Education, Youth and Sports of the Czech Republic. Part of the work was carried out with the support of a Growth Facility (BC Core Facilities; IPMB BC CAS). X. laevis oocytes were kindly provided by C. Korbmacher on a regular basis (FAU Erlangen-Nürnberg). MF received support from the European Research Council (Grant 480 No. 101125499). We acknowledge the core facility LMH, the BC CAS supported by the MEYS CR (LM 2023050 Czech-BioImaging). DO received support from the Czech Science Foundation grant Nr. 24-12107S. LCMS analyses were performed in Laboratory of Mass Spectrometry at Biocev research center; Faculty of Science, Charles University.

## Supplementary Movies

**Supplementary Movie 1**

Co-localization of ARO2-GFP and CNGC14-mSCARLET under their native promoters in *A.thaliana* root elongation zone.

**Supplementary Movie 2**

Kinetics of the root gravitropic bending of *aro2/4* and *aro2/3/4* mutants. Note that ARO3 is not expressed in the root elongation zone.

**Supplementary Movie 3**

pH profile of the *aro2/3/4* and WT roots during the course of root gravitropic bending. Magenta represents acidic shift; green represents alkaline shift.

**Supplementary Movie 4**

Microfluidic setup showing GCaMP (green) in *aro2/3/4* and WT, incubated side by side in a single flow channel for identical conditions. Blue hue represents treatment with 10nm IAA.

**Supplementary Movie 5**

Plant leaves exposed to drought stress show calcium transients in stomata (asterisks) visualized by GcaMP fluorescence.

---

**Figure S1.**
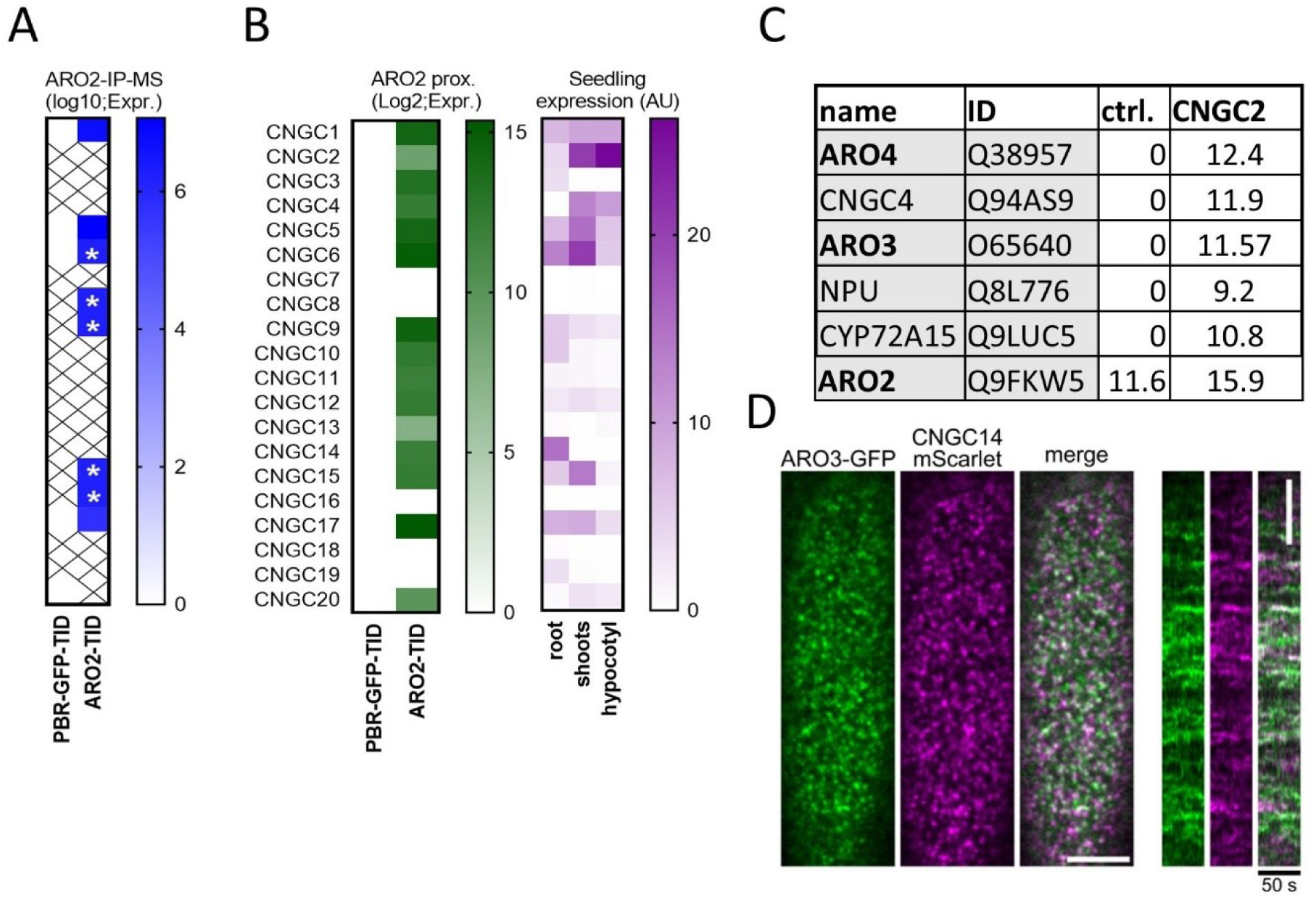
AROs interact with CNGCs. **(A)** IP-MS results on the ARO2-tID-HA compared to PBR-GFP-tID-HA. Asterisks depict ambiguous peptides **(B)** Summary of all the CNGCs identified in the ARO2 proxiome in comparison with their expression levels in the seedlings^28^ **(C)** Top five most enriched proteins in CNGC2 proxiome. ARO2 enrichment was lower due to signal in neg. control. Values represent mean binary logarithms of protein intensity. The experiment was done in triplicate. **(D)** Co-localization of ARO3-GFP and CNGC14-mSC in the differentiating trichoblast. Right – line intensity profile over 50 seconds. Bar=5µm.

**Supplementary figure S2.**
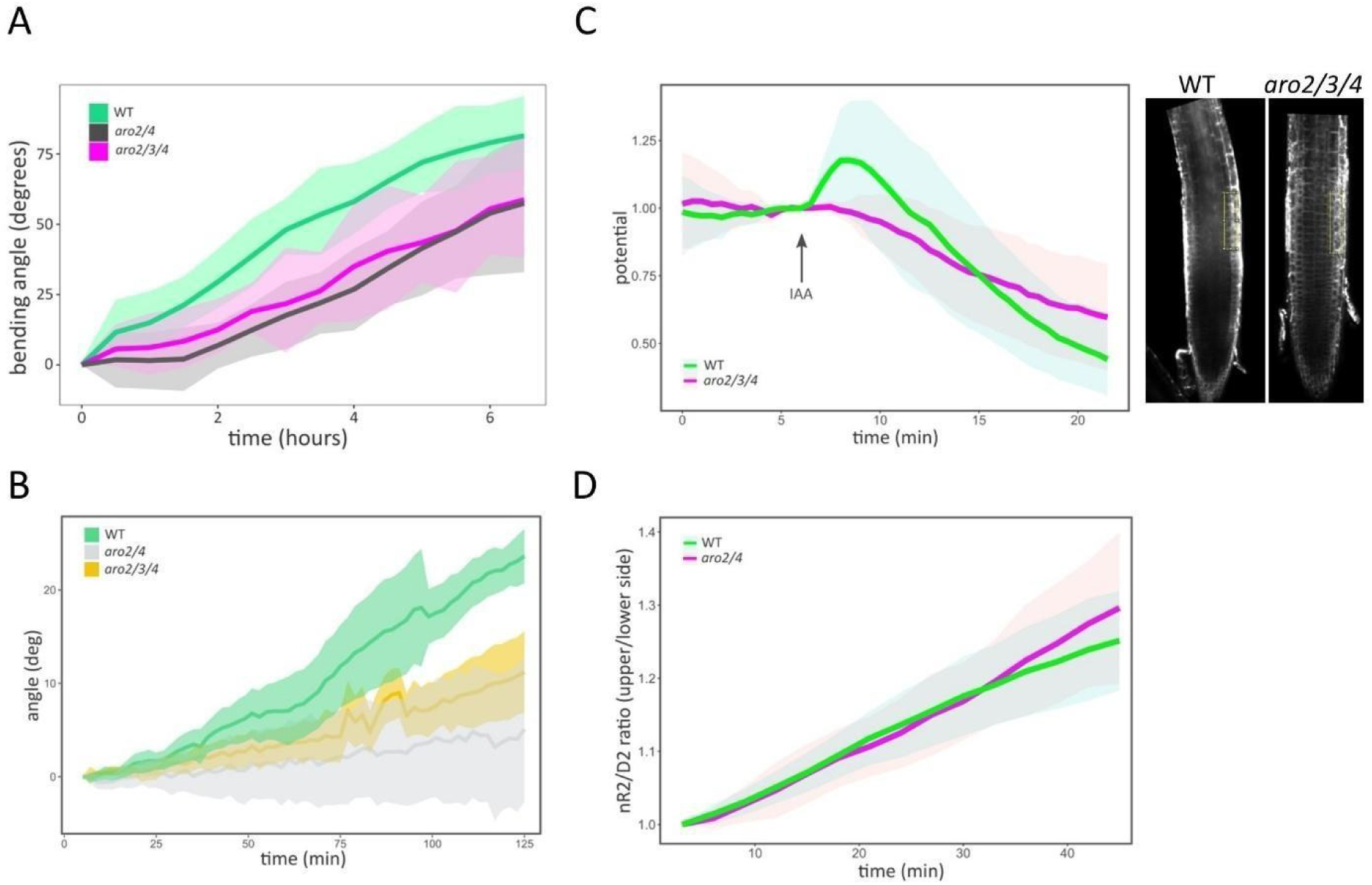
AROs are involved in auxin responses downstream of auxin transport and upstream of membrane depolarization. **(A)** Gravitropic bending of the *aro2/3/4* and *aro2/4* mutants grown within the agar media. Average of 8 plants. Coloured areas represent SD. Multiple similar replicas were generated. **(B)** Gravitropic bending of the *aro2/3/4* and *aro2/4* mutants grown on the media surface, with high temporal resolution. Average of 5 plants per genotype. Coloured areas = SD. **(C)** *aro2/3/4* mutants fail to depolarize the PM upon 20nm IAA treatment using DISBAC_2_(3) voltage sensor. 5 plants per genotype were used. Coloured areas = SD. Pictures on the right depict the area which was evaluated. **(D)** Transcriptional responses to auxin, evaluated using R2D2 ratio upon gravitropic bending, remain unaffected in *aro2/4* mutants. Average of >8 roots per sample.

**Supplementary figure S3.**
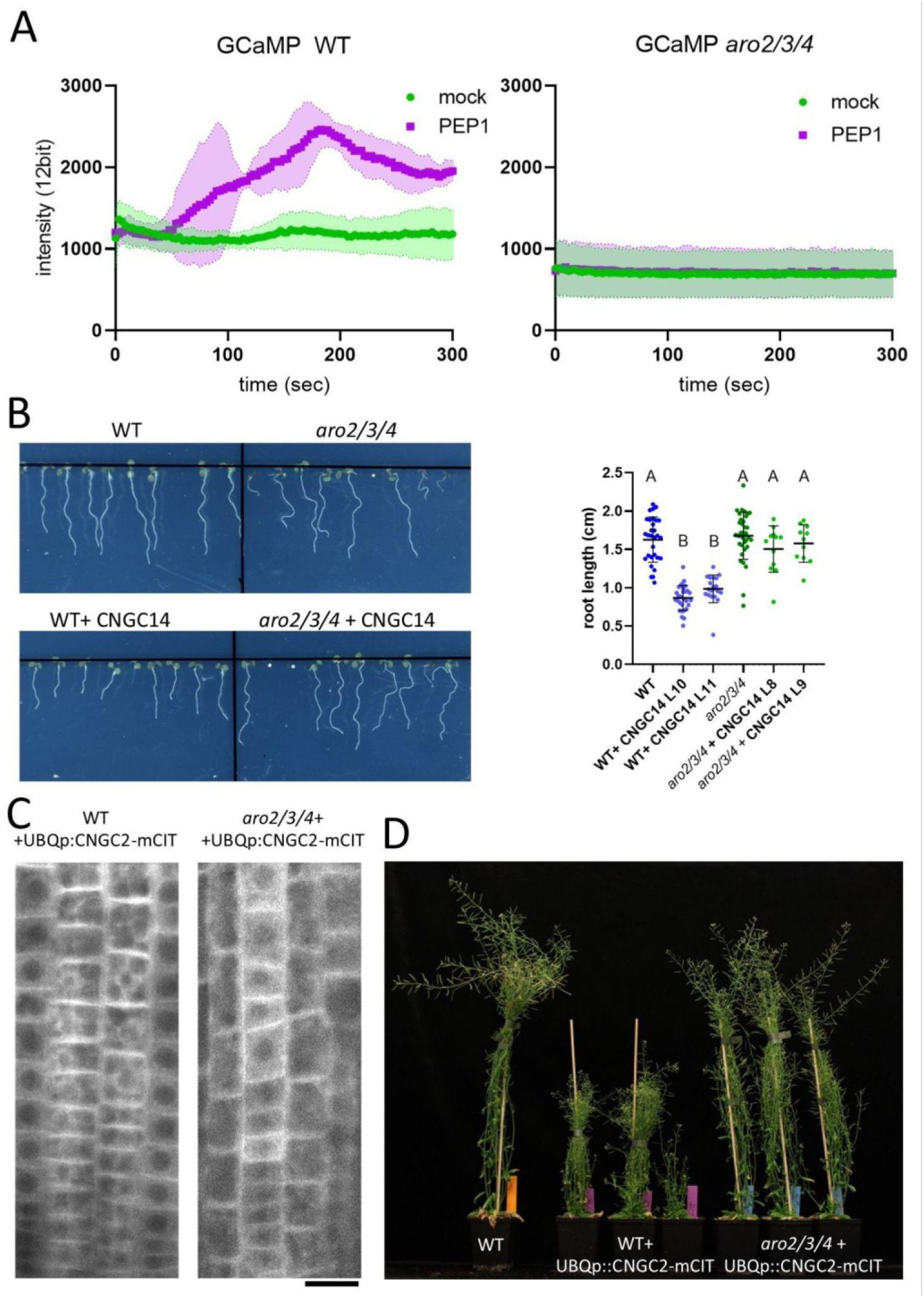
*aro2/3/4* mutants are resistant to CNGC overexpression. **(A)** Calcium transient triggered by the 1µM PEP1 peptide in the WT and *aro2/3/4* mutants. The experiment was repeated 3 times. **(B)** Left **-** Representative 7-day-old seedlings of WT and *aro2/3/4* expressing CNGC14p:CNGC14-GFP. Right- quantification of the root length. Addition of transgenic CNGC14 results in shorter root in WT, but not in *aro2/3/4* mutants. **(C)** Representative pictures of the plants overexpressing UBQp:CNGC2-mCIT construct. Bar = 20μm. **(D)** CNGC2 overexpression manifests as smaller WT plants, while *aro2/3/4* plants display normal size. 3 plants display 3 independent transgenic lines.

**Supplementary figure S4.**
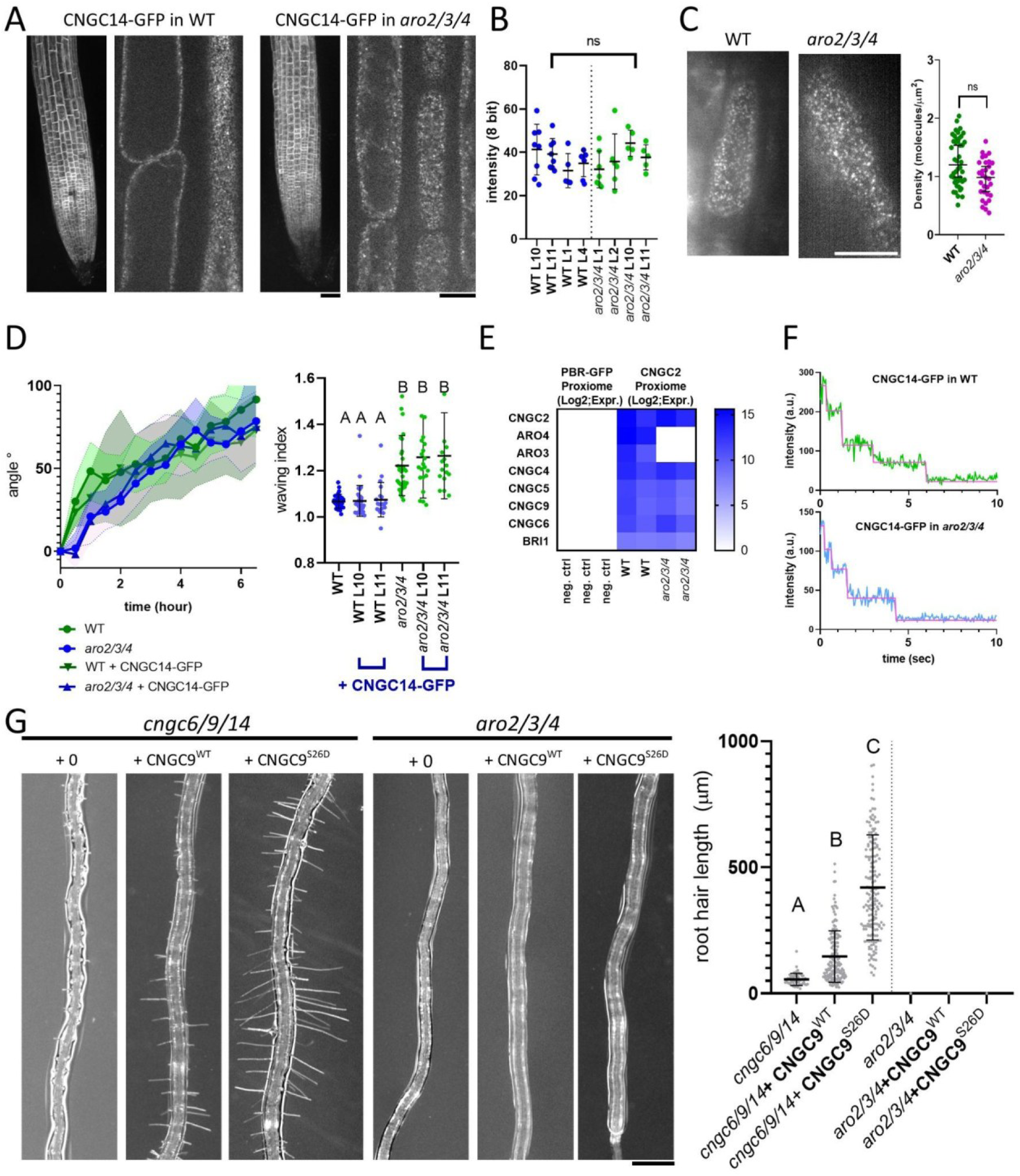
CNGCs localize and multimerize normally in *aro2/3/4* mutant background. **(A)** CNGC14-GFP under its native promoter displays similar localization in *aro2/3/4* mutant and WT. Left – overview of the whole root. Right – cortical section of the PM displaying similar punctate pattern. Bars=50µm (left), 10µm (right) **(B)** CNGC14–GFP displays comparable signal intensity among different T-DNA lines. **(C)** CNGC14-GFP displays comparable amount of PM puncta at the plasma membrane using TIRF microscopy. Bar=10µm **(D)** *aro2/3/4* mutants and WT plants expressing similar amounts of CNGC14-GFP still retain the gravitropic bending phenotype and root waving phenotypes. **(E)** In the CNGC2 proxiome, CNGC4 and other CNGCs, as well as BRI1 are biotinylated equally regardless of presence of AROs. CNGC2 and AROs are included as an internal control. CNGC4 was the second most differentially biotinylated protein after ARO4. **(F)** Representative images from the SMPB experiments showing that also in *aro2/3/4* mutant lines, 4 bleaching steps (CNGC14 tetramers) can be found. >10 similar replicates were generated for both genotypes. **(G)** left**-** Representative images of the *cngc6/9/14* and *aro2/3/4* transformed with CNGC9 WT and phospho-mimetic versions. Right - Quantification of D. 5 Average of 10 different T1 plants. Letters denote statistical significance (One-way Anova p<0.0001).

**Supplementary figure S5.**
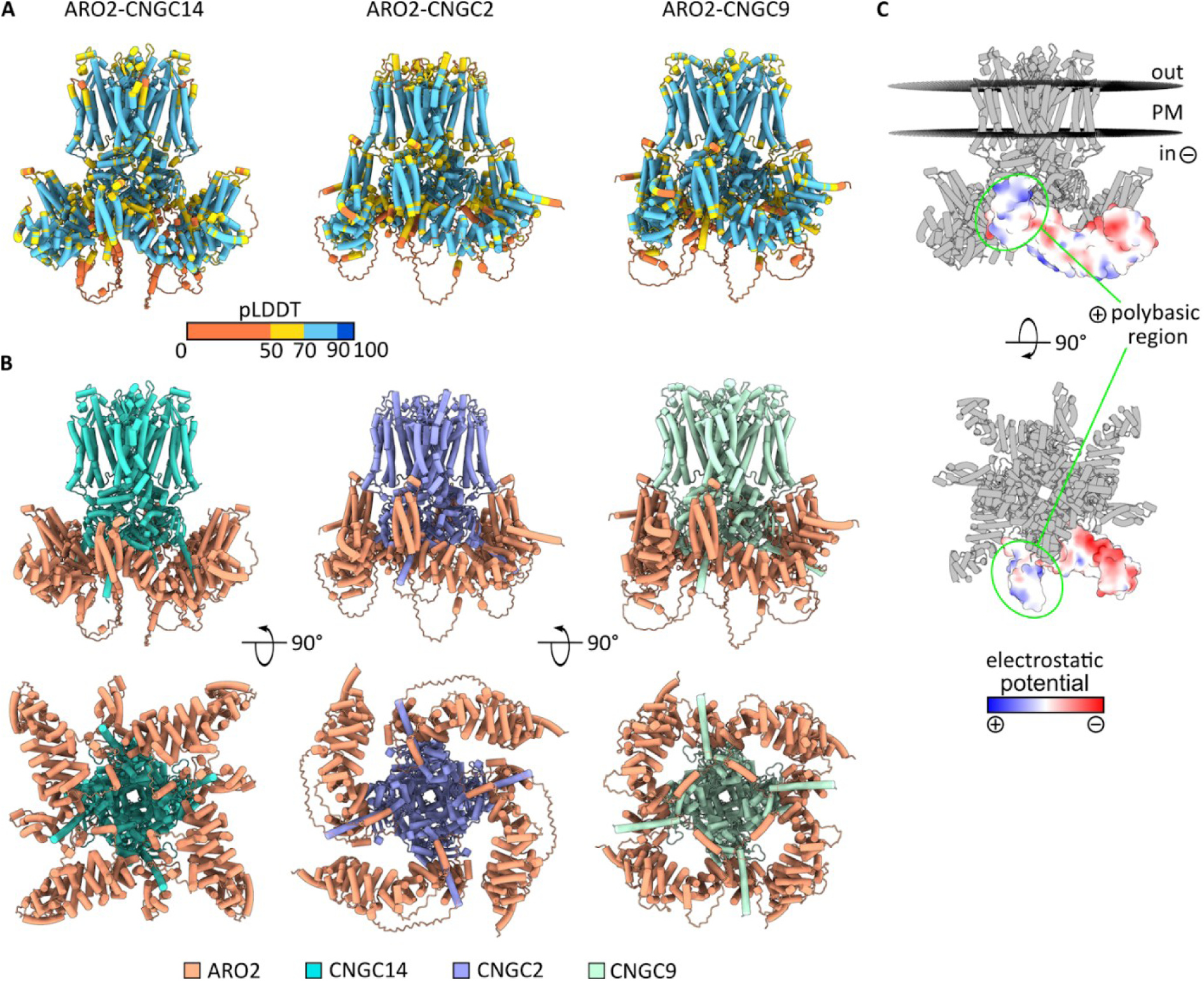
Structural models of ARO2-CNGC14, ARO2-CNGC2 and ARO2-CNGC9. **(A)** AlphaFold3 predictions of ARO2-CNGC14, ARO2-CNGC2 and ARO2-CNGC9 complexes colored according to pLDDT score. **(B)** AlphaFold3 predictions of the ARO2–CNGC14, ARO2–CNGC2, and ARO2–CNGC9 complexes, with each subunit shown in a distinct color. **(C)** The PPM3.0 positioning of the ARO2-CNGC14 complex in the membrane. Electrostatic potential mapped on one of the ARO2 subunits shows the N-terminal polybasic region, previously described to mediate the interaction with anionic phospholipids of the PM^25^ is oriented towards the membrane and thus enabling its interaction with negatively charged phospholipids.

**Supplementary Figure S6.**
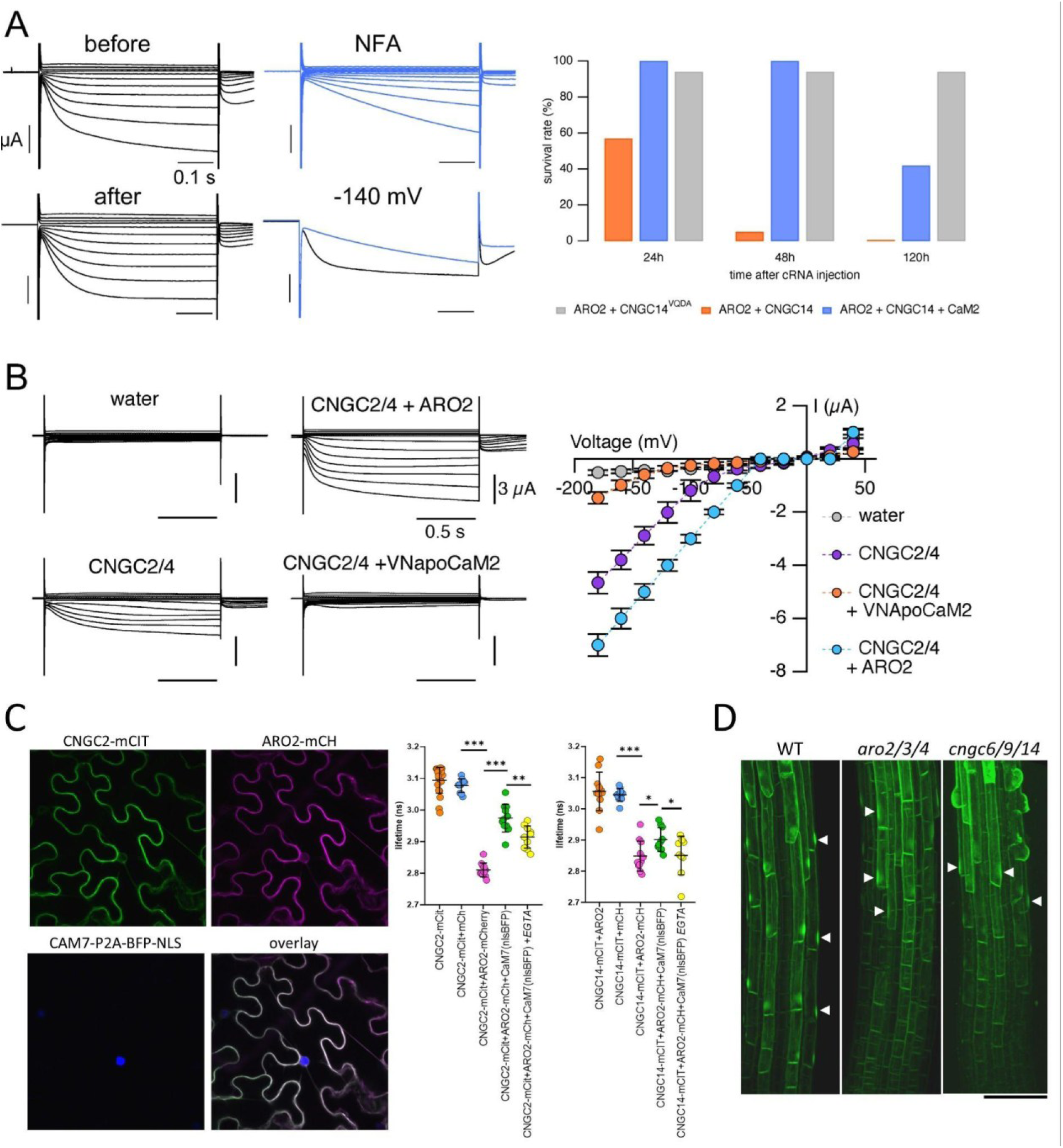
Regulation of CNGCs by ARO2 and Ca^2+^-CaM2 variants. **(A)** CNGC14 activity elicits calcium currents amplified by endogenous Ca^2+^-dependent chloride channel currents and co-expression of ARO. *Left:* Currents of a representative *Xenopus* oocyte expressing CNGC14 in a solution containing 30 mM CaCl_2_ before and after application of the chloride channel inhibitor niflumic acid (NFA) at 300 µM, and after wash out. *Right:* Cytotoxicity of CNGC14 and ARO2 in *Xenopus* oocytes is dependent on binding to the IQ motif. Survival rates of injected oocytes 24, 48 and 120 hours after co-injection of CNGC14 and ARO2 (n = 21), CNGC14, ARO2 and CaM2 (n = 19), or the IQ motif mutant CNGC14VQ/DA and ARO2 (n = 18). Dead oocytes were removed and the ND96 solution replaced after 24 and 48 hours. **(B)** ARO2 activates CNGC2/4 heterotetrameric channels. Current responses (left) and corresponding current-voltage relations (right) of oocytes injected with water (n=3), or co-injected with CNGC2 and CNGC4, together with water (n=6), VN-apoCaM2 (n=6) or ARO2 (n=8). Mean currents ± SEM are shown, resulting from CNGC2/4 activity amplified by Ca^2+^-activated endogenous chloride currents. Currents were recorded in a solution containing 30 mM CaCl_2_. **(C)** Cam7 and ARO2 compete for CNGC interaction. Left- representative image showing the setup, where CAM7 was introduced with cleavable BFP-NLS. Right – quantification of the CNGC2- and CNGC14-ARO interactions by FRET-FLIM analysis. Asterisks denote statistical significance (T-test *p<0.05; **p<0.005;***p<0.0005). **(D)** Distribution of GFP-ROP2, a marker of the root hair initiation. Root hair initiation sites (arrows) are gradually diffusing in both *aro2/3/4* and *cngc6/9/14* mutants, arguing for polarity defects of aro2/3/4 being downstream of Ca^2+^ signaling. Bar=100µm.

## Notes

### Competing Interest Statement

The authors have declared no competing interest.

### Summary of Updates

We added new ARO and CNGC proxiome data and improved electrophysiology data. We address phosphorylation and calmodulin crostalk with ARO mediated regulation of the CNGC activity.

